# An expanded proteomic survey of the human parasite *Leishmania major* focusing on changes in null mutants of the Golgi GDP-Mannose/Fucose/Arabino*pyranose* transporter *LPG2* or the mitochondrial fucosyltransferase *FUT1*

**DOI:** 10.1101/2022.10.29.514353

**Authors:** Gloria Polanco, Nichollas E. Scott, Lon F. Lye, Stephen M. Beverley

## Abstract

The trypanosomatid protozoan parasite *Leishmania* has a significant impact on human health globally. Understanding the pathways associated with virulence within this significant pathogen is critical for identifying novel vaccination and chemotherapy targets. Within this study we leverage an ultradeep proteomic approach to improve our understanding of two virulence associated genes in *Leishmania*, encoding the Golgi Mannose/Arabino*pyranose*/Fucose nucleotide-sugar transporter *LPG2*, and the mitochondrial fucosyltransferase *FUT1*. Using deep peptide fractionation followed by complementary fragmentation approaches with higher energy collisional dissociation (HCD) and Electron-transfer dissociation (ETD) allowed the identification of over 6500 proteins, nearly doubling the experimentally known *Leishmania major* proteome. This deep proteomic analysis revealed significant quantitative differences in both *Δlpg2^-^* and *Δfut1^s^* mutants with *FUT1*-dependent changes linked to marked alterations within mitochondrial associated proteins while *LPG2*-dependent changes impacted many pathways including the secretory pathway. While the FUT1 enzyme has been shown to fucosylate peptides *in vitro*, no evidence for protein fucosylation was identified within our ultradeep analysis nor did we observe fucosylated glycans within *Leishmania* glycopeptides isolated using HILIC enrichment. Combined this work provides a critical resource for the community on the observable *Leishmania* proteome as well as highlights phenotypic changes associated with *LPG2* or *FUT1* ablation which may guide the development of future therapeutics.

**Importance:** *Leishmania* is a widespread trypanosomatid protozoan parasite of humans with ∼12 million cases ranging from mild to fatal, and hundreds of millions asymptomatically infected. This work advances knowledge of the experimental proteome by nearly 2 fold, to more than 6500 proteins a great resource to investigators seeking to decode how this parasite is transmitted and causes disease, and new targets for therapeutic intervention. The ultradeep proteomics approach identified potential proteins underlying the ‘persistence without pathology’ phenotype of deletion mutants of the Golgi nucleotide transporter LPG2, showing many alterations and several candidates. Studies of a rare deletion mutant of the mitochondrial fucosyltransferase FUT1 revealed changes underlying its strong mitochondrial dysfunction, but did not reveal examples of fucosylation of either peptides or N-glycans. This suggests this vital protein’s elusive target(s) may be more complex than the methods used could detect, or may not be a protein, perhaps another glycoconjugate or glycolipid.

## Introduction

Leishmaniasis is a devastating parasitic disease caused by species of the trypanosomatid protozoan parasite genus *Leishmania*. Currently, there are over 12 million cases, with 1.7 billion people at risk of infection world-wide, and estimates of asymptomatic infections ranging as high as >300 million (1–4). Clinical manifestations include self-healing localized or diffuse cutaneous lesions (cutaneous leishmaniasis), destruction of the nasopharyngeal mucosa (mucocutaneous leishmaniasis), or enlargement of spleen or liver which can lead to death (visceral leishmaniasis) (3). *Leishmania* parasites are transmitted to mammalian hosts via the bite of phlebotomine sand flies and undergo a series of transformations throughout its life cycle, predominantly as promastigotes (insect stage) and amastigotes (intracellular in mammalian host) (3, 5). Powerful genetic tools have uncovered a number of loci essential to the completion of the parasites life cycle, and/or potentially suitable as targets for chemotherapy (6–10). Here, we focus on two mutants relevant to such efforts, whose properties are depicted in Figure 1.

**Figure 1.**
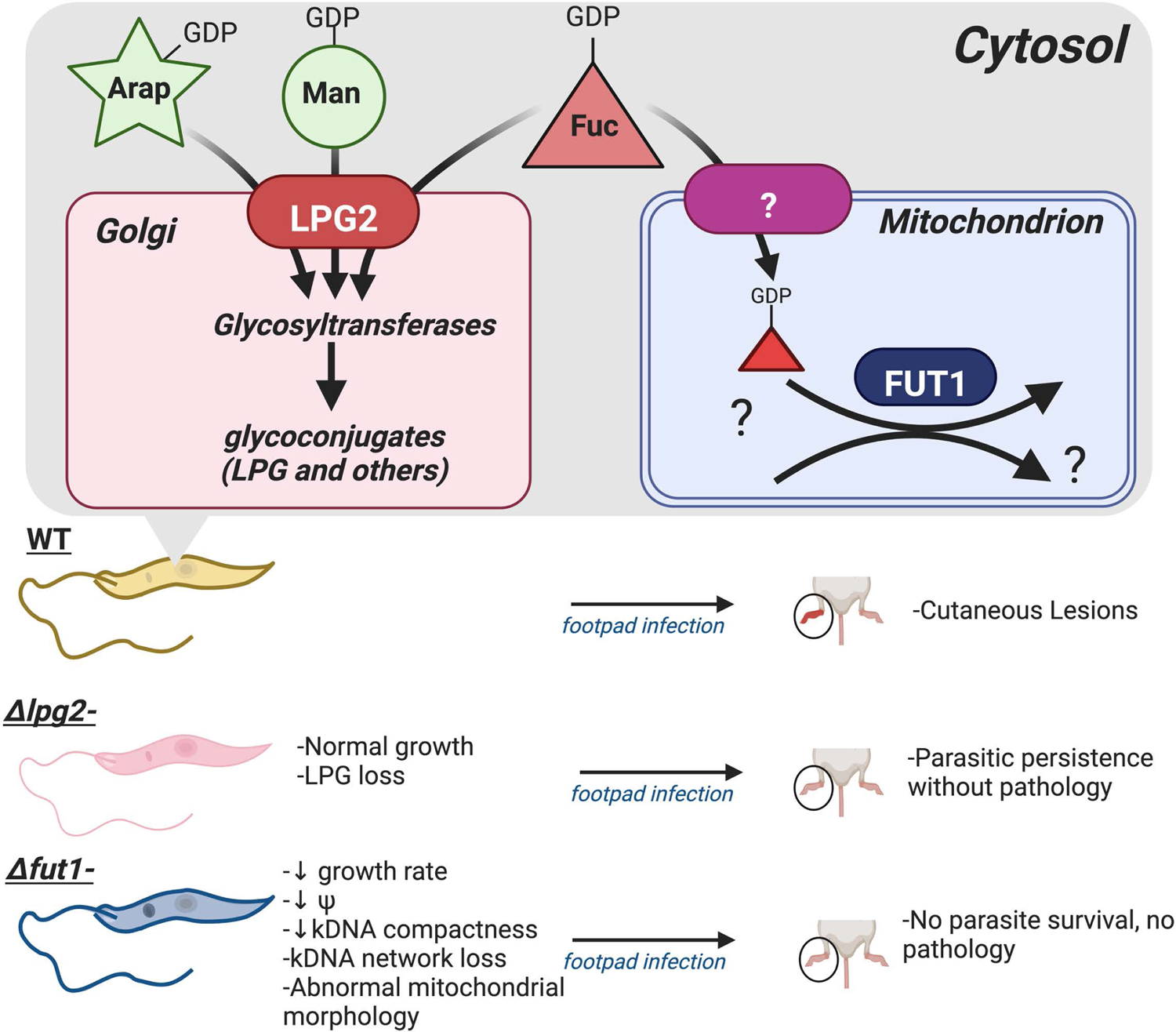
Over view of glycosylation pathways relevant to *LPG2* or *FUT1* biology and deletion mutants*. LPG2* encodes is a Golgi GDP-sugar transporter, essential for LPG synthesis. Mutant *Δlpg2^-^* parasites grow comparable to WT parasites in culture, but do not produce pathology. FUT1 encodes a fucosyltransferase located in the parasite mitochondria, whose substrates remain unknown. Null mutant *Δfut1**^s^***parasites exhibit severe defects including significantly decreased growth rate, abnormal mitochondrial morphology, decreased mitochondrial membrane potential (Ψ), loss in kDNA compactness or the kDNA network altogether. Abbreviations and symbols are: arabino*pyranose* (Ara*p*): filled green star; mannose (Man): filled circle; fucose (Fuc): filled triangle; LPG: lipophosphoglycan.

*Leishmania* parasites are completely coated in a dense glycocalyx largely containing glycoconjugate lipophosphoglycan (LPG), composed of 1) a 1-O-alkyl-2-lyso-phosohotidylinositol lipid anchor, 2) a heptasaccharide core, 3) a phosphoglycan polymer consisting of 15-30 [Galβ1,4Manα1-PO_4_] repeating units (phosphoglycan or PG repeats), often bearing side chain sugars such as galactose in *L. major,* and 4) a small oligosaccharide cap (11–13). Biochemical and genetic studies of *L. major Δlpg1*^-^ mutants specifically lacking LPG have implicated it in many key steps of the parasite infectious cycle: in the sand fly vector, as well in the establishment of infection within the mammalian host following the sand fly bite (14–18). However, the amastigote stage lacks significant levels of LPG, and those few *Δlpg1*^-^ parasites able to survive the initial host response are thereafter highly virulent (17). In contrast, deletion of *LPG2* encoding a Golgi nucleotide sugar transporter (NST; described below) resulted in loss of LPG along with related glycoconjugates including proteophosphoglycans, which are normally expressed in amastigotes (19). While many *Δlpg2^-^* phenotypes in both sand flies and establishment of mammalian infection mirrored those of *Δlpg1*^-^, *Δlpg2^-^* infections of susceptible mice show no overt disease and long-term persistence of a small number of parasites typical of long-term asymptomatic infections (19). This ‘persistence without pathology’ phenotype induced protective immunity in a manner similar to that seen in natural healed infections (20).

While the attenuated *Δlpg2^-^* phenotype had been attributed solely to the loss of LPG and related glycoconjugates (19), this view was challenged by studies of an analogous LPG-deficient mutant obtained by genetic deletion of the partially redundant *LPG5A* and *LPG5B* genes encoding the Golgi UDP-Galactose transporters (21). These parasites resemble *Δlpg2^-^* in lacking LPG and related PG-bearing glycoconjugates, but unlike *Δlpg2^-^*, retained pathology following inoculation into susceptible mice similar to that seen in *Δlpg1*^-^ mutants lacking only LPG (22). This suggested the possibility of *LPG2*-dependent “off-LPG/phosphoglycan” effects, identification of which could shed new insight on both the persistence without pathology phenotype of *Δlpg2^-^*, its ability to serve as a live vaccine line, and its tendency to revert towards virulence via second-site events (20, 23, 24).

The second mutant studied arises from finding that LPG2 was the first example of a multi-specific NST, able to transport GDP-L-fucose as well as GDP-D-Ara*pyranose* (25). While the roles of GDP-Man in phosphoglycan repeat synthesis and D-Ara*p* as a ‘capping’ sugar able to block LPG interaction with the sand fly epithelium had been thoroughly studied (26, 27), GDP-Fuc transport was enigmatic. Although *L. major* bore low levels of GDP-Fucose mediated by the closely related bifunctional salvage enzymes AFKP80 and FKP40 (28, 29), convincing evidence for fucoconjugates has been hard to find. However, deletion of the two salvage enzymes could not be achieved unless ‘metabolic complementation’ of GDP-fucose was engineered through expression of the trypanosome *de novo* GDP-fucose synthetic pathway (29). Reasoning this implied the existence of an essential fucosyltransferase (FUT)., we identified 5 candidate fucosyltransferases, four of which appeared to be targeted as expected to the parasite secretory pathway. *SCA1* and *SCA2* function in D-Ara*p* modifications of LPG (26) while null mutants of *SCAL* and *FUT2* showed little phenotype in culture (30).

The fifth candidate *FUT1*, however, was important for parasite survival and potentially the key player in the GDP-Fucose requirement. Unlike the other candidates, *FUT1* was found throughout all trypanosomatid species but not in other organisms (30, 31). Unexpectedly, FUT1 targeted to the parasite mitochondrion, as was its homolog TbFUT1 from *Trypanosoma brucei*, and both the mitochondrial localization and fucosyltransferase activity were essential (30, 31). While impossible to knockout by conventional approaches, through plasmid shuffling and examining more than 1000 events, a single, rare *Leishmania* Δ*fut1* deletion segregant (Δ*fut1^s^*) was obtained (30). This mutant line displayed severe growth and mitochondrial defects, which were rescued by Lmj*FUT1* or Tb*FUT1* re-expression (30). While FUT1s described in other organisms are able to fucosylate a variety of glycan substrates (32, 33), recombinant *L. major* FUT1 was unexpectedly able to fucosylate both glycan and peptide substrates *in vitro* (30). Thus far the native acceptor in neither *Leishmania* nor trypanosomes has been identified despite considerable effort in both species (30, 31).

Here, we generated high coverage proteomes of WT, *Δlpg2^-^* and *Δfut1^s^ Leishmania major,* identifying over 6500 proteins representing nearly 80% of the predicted proteome, expanding the previously known experimental *L. major* proteome by nearly 2-fold. Differential proteomic analysis showed numerous differences in the *Δlpg2^-^*or *Δfut1^s^* parasites, including some impacted in both. Despite the depth of this analysis, we failed to identify any fucose-bearing proteins across these proteomes. Furthermore while glycopeptide enrichment analysis of WT samples also identified numerous instances of N-linked glycopeptides, no O-fucosylation or fucosylated N-linked glycans were observed. Taken together this work highlights the challenges of identifying FUT1 targets and that alternative approaches will be required in future studies.

## Results

### Acquisition of a high coverage of *L. major* proteome

We examined the WT Fn strain of *L. major*, whose genome is one of the best characterized references for *Leishmania* sp., as well as *L. major* Fn *Δfut1^s^* (30) and a newly created Fn *Δlpg2^-^* mutant. CRISPR/Cas9 mutagenesis was used to readily generate homozygous ORF replacements (Fig. S1), and one Fn *Δlpg2^-^* clonal line (c14.2) was chosen for further study. Preliminary studies showed it lacks LPG by agglutination tests, does not give lesion pathology when inoculated into susceptible mice, but persists at low levels, similar to the previously described strain *L major* LV39 clone 5 *Δlpg2^-^*mutant (19). Previous studies showed that restoration of *LPG2* expression restores all WT phenotypes tested in both the Fn and LV39 clone 5 backgrounds (19, 34),

For each line, four replicate cultures were initiated and harvested in logarithmic growth phase. Parasite lysates were digested with trypsin and separated into 12 concatenated fractions by basic reverse phase C18 chromatography (35), and then individually separated and analyzed by LC-MS/MS. To ensure the ability to localize any potential glycosylation events, precursors were subjected to both higher energy collisional dissociation (HCD) and electron-transfer dissociation (ETD) fragmentation with spectra searched against a database consisting of predicted *L. major* proteome. Protein matches meeting a 0.01 FDR cutoff, excluding contaminants, reverse decoys and those only identified by site were retained for an initial analysis of the coverage (Table S1).

The number of proteins detected was similar among WT, *Δlpg2^-^*, and *Δfut1^s^* (6208, 6102, and 5934, respectively), collectively totaling 6,744 (Table S1) and representing nearly 80% of the predicted *L. major* proteome (8307 or 8038 in TriTrypDB or UniProt respectively). In comparisons with two previous proteomic studies of *L. major* (36, 37) 3,484 proteins identified had no prior MS-based evidence, while 3,260 had prior MS-based evidence, and 308 detected previously did not appear in our dataset (Fig. 2A). Thus, this work increased experimental coverage of the *Leishmania* proteome by nearly 2-fold.

**Figure 2.**
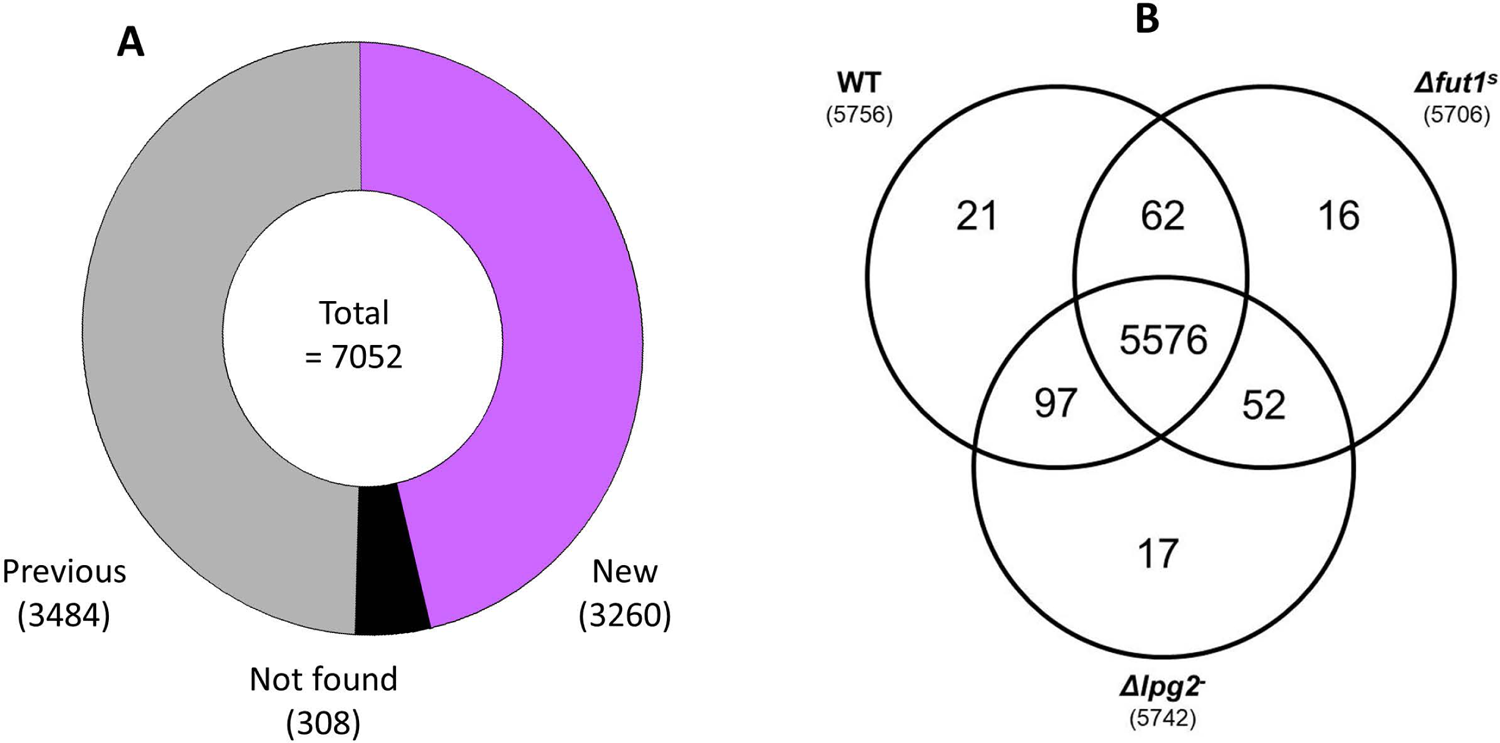
<u>A. Comparison of proteins identified in this work with previous studies.</u> The combined dataset (Table S1) from all lines and replicas was compared to *L. major* proteins with MS based evidence annotated in TriTrypDB (www.TryTryp.org). 3484 proteins identified here showed previous MS evidence, 3260 proteins had no prior MS-based reports, and and 308 proteins with previous MS-based evidence did not appear in our datasets. <u>B.</u> Distribution of identified proteins amongst WT, *Δfut1**^s^***, and *Δlpg2^-^ L. major*. The total proteome (Table S1) was further parsed by considering only proteins identified in at least two of the four biological replicates, in one or more parasite line, yielding a total of 5841 (Table S3). The venn diagram displays the overlap of proteins amongst the three lines.

No predicted mitochondrion-encoded proteins were identified, despite the inclusion of many of them in our predicted proteome database (described in the Methods). Often these are omitted from proteomic computational analysis completely, and due to their hydrophobicity, they are often not detected by standard or general approaches similar to those used here (38–40).

### Protein glycosylation and absence of detectable fucosylation

We showed previously that recombinant FUT1 is able to modify both glycan and peptide substrates *in vitro* (30) and the expanded proteome provided an opportunity to search for evidence of *in vivo* fucosylation. To allow the localization of fucosylation events ETD MS fragmentation was used as O-fucosylated peptides are poorly localized with collision based approaches (41). We searched first for peptide-O-fucose by considering deoxyhexose (dhex) modifications of serine or threonine residues yet despite our proteome depth and the ability of our group to identify glycosylation events from deep proteome dataset (42, 43) no high confidence O-fucosylation events were observed. Moreover, we were specifically unable to detect HSP70 or HSP60 peptide O-fucose, inferred previously in *Leishmania donovani* (44), peptides of which were fucosylated by recombinant FUT1 *in vitro* (30). Open searching (45) was also undertaken using MSFragger (46) which also failed to reveal any putative fucose containing glycopeptides from our deep proteome analysis.

To further explore if the diversity of glycans which may exist within *L. major* and we surveyed glycopeptides derived from *L. major* using zwitterionic–hydrophilic interaction liquid chromatography (ZIC-HILIC) (47) coupled to LC-MS analysis. Using Open searching with MSFragger (46) multiple unique glycoforms were observed, majority of which corresponded to N-linked glycoforms. While short O-linked glycopeptides can be enriched with this approach (48) our observation of predominately N-linked glycopeptide support that these escaped detection from *L. major* using ZIC-HILIC based enrichment. We identified 65 glycopeptides bearing a variety of high-mannose N-linked glycans as reported previously in *Leishmania* (49–52). The glycopeptides mapped to 49 different proteins, with 14 showing multiple sites and/or glyco-heterogeneity; 24 were predicted as hypothetical proteins of unknown function. The annotated glycosylated proteins are summarized in Table 1 and include many described previously, such as gp63/leishmanialysin, nucleotidases, and phosphatases (50), or ones likely to be so due so do the presence of targeting motifs for the secretory pathway such as *FUT2* (30). While these finding highlight that multiple N-linked glycopeptides could be recovered from *L. major,* no fucosylated N-linked glycans were observed (Table S2). Within this dataset, three additional proteins absent from the total proteome were identified, bringing the collective total identified to 6747. The three were all hypothetical proteins, two without prior MS-based evidence (LmjF.08.0350, LmjF.09.1330) and one previously reported (LmjF.33.1035) (36). Both LmjF.08.0350 and LmjF.09.1330 have a detected mass shift of approximately 1380.5 Da, while LmjF.33.1035 presented a mass shift of 2803.3 Da. Future work will be required to confirm these preliminary assignments and the potential identity of the attached modifications.

**Table 1:** A first-pass *L. major* glycoproteome from MS analysis of HILIC-enriched glycopeptides

| **Protein Annotation** | **ID** |
| --- | --- |
| ATP pyrophosphate-lyase | LmjF.17.0200 |
| $\beta$ galactofuranosyl transferase (*LPG1*) | LmjF.25.0010 |
| C-8 sterol isomerase-like protein | LmjF.29.2140 |
| Cathepsin L-like protease | LmjF.08.1060 |
| Chitinase | LmjF.16.0790 |
| D-alanyl-glycyl endopeptidase-like protein | LmjF.33.2850 |
| Dolichyl-diphosphooligosaccharide--protein glycotransferase | LmjF.35.1130 |
| GPI ethanolamine phosphate transferase 3 | LmjF.24.0340 |
| Hypothetical predicted multi-pass transmembrane protein | LmjF.24.0700 |
| Leishmanolysin (GP63) | LmjF.10.0465 |
| Phosphoglycan $\beta$ 1,3 galactosyltransferase 3 (*SCG3*) | LmjF.02.0010 |
| Putative 3'-nucleotidase/nuclease | LmjF.12.0400 |
| Putative 3'-nucleotidase/nuclease | LmjF.31.2310 |
| Putative amastin-like surface protein | LmjF.34.1080 |
| Putative β-fructofuranosidase | LmjF.04.0310 |
| Putative DNAJ domain protein | LmjF.24.0520 |
| Putative extracellular receptor | LmjF.19.0640 |
| Putative glycosyl transferase (*FUT2*) | LmjF.02.0330 |
| Putative phospholipid:diacylglycerol acyltransferase | LmjF.09.1040 |
| Putative surface protein amastin | LmjF.30.0860 |
| Signal recognition particle receptor subunit $\beta$ | LmjF.33.2620 |
| Signal sequence receptor subunit alpha | LmjF.22.0260 |
| Stealth_CR3 domain-containing protein | LmjF.16.1010 |
| Transmembrane 9 superfamily member | LmjF.34.3660 |
| Uncharacterized protein L7845.03 (Thioredoxin) | LmjF.35.1330 |

The table shows a list of annotated proteins identified in the glycoproteomic analysis in this work. The gene IDs and the protein annotations are taken from the TriTrypDB (www.tritrypdb.org). Another 24 hypothetical proteins lacking annotation were also identified.

### WT and Mutant Cell Lines Display a Similar Qualitative Proteome

To compare the impact of *LPG2* or *FUT1* loss, we first created a ‘high confidence’ data set by parsing the total proteome (Table S1) for proteins detected in two or more biological replicates, in at least one cell line. This retained 5841 proteins (Table S3), of which 95% (5576) occurred in all three lines (Fig. 2B), evidence of high congruency. Of the remainder, 16 - 21 were unique to one, and 62 - 97 were shared by two lines (Fig. 2B). As expected, the LPG2 protein was detected in all WT and *Δfut1^s^*lysates but not in *Δlpg2^-^* replicas. FUT1 peptides, on the other hand, was detected only in one WT biological replicate, perhaps due to low abundance (53).

### Mitochondrial proteome and Gene Ontology

As FUT1 is targeted to the mitochondrion where its loss results in numerous mitochondrial abnormalities (30), we queried the high confidence proteomic data set for nuclearly-encoded mitochondrial proteins present in the trypanosomatid predicted mitochondrial protein database MiNT (54). Of 1559 mitochondrial *L. major* proteins in the MiNT database (approximately 19% of total proteome), 1268 (81%) were identified (Fig. 3A; Table S3), with WT having a slightly higher number of identified proteins (1241), followed by *Δlpg2^-^* (1239) and *Δfut1^s^* (1226; Fig. 3B). Quantitative changes were apparent (as discussed in a later section), and no mitochondrial (maxicircle) encoded proteins were identified as discussed earlier.

**Figure 3.**
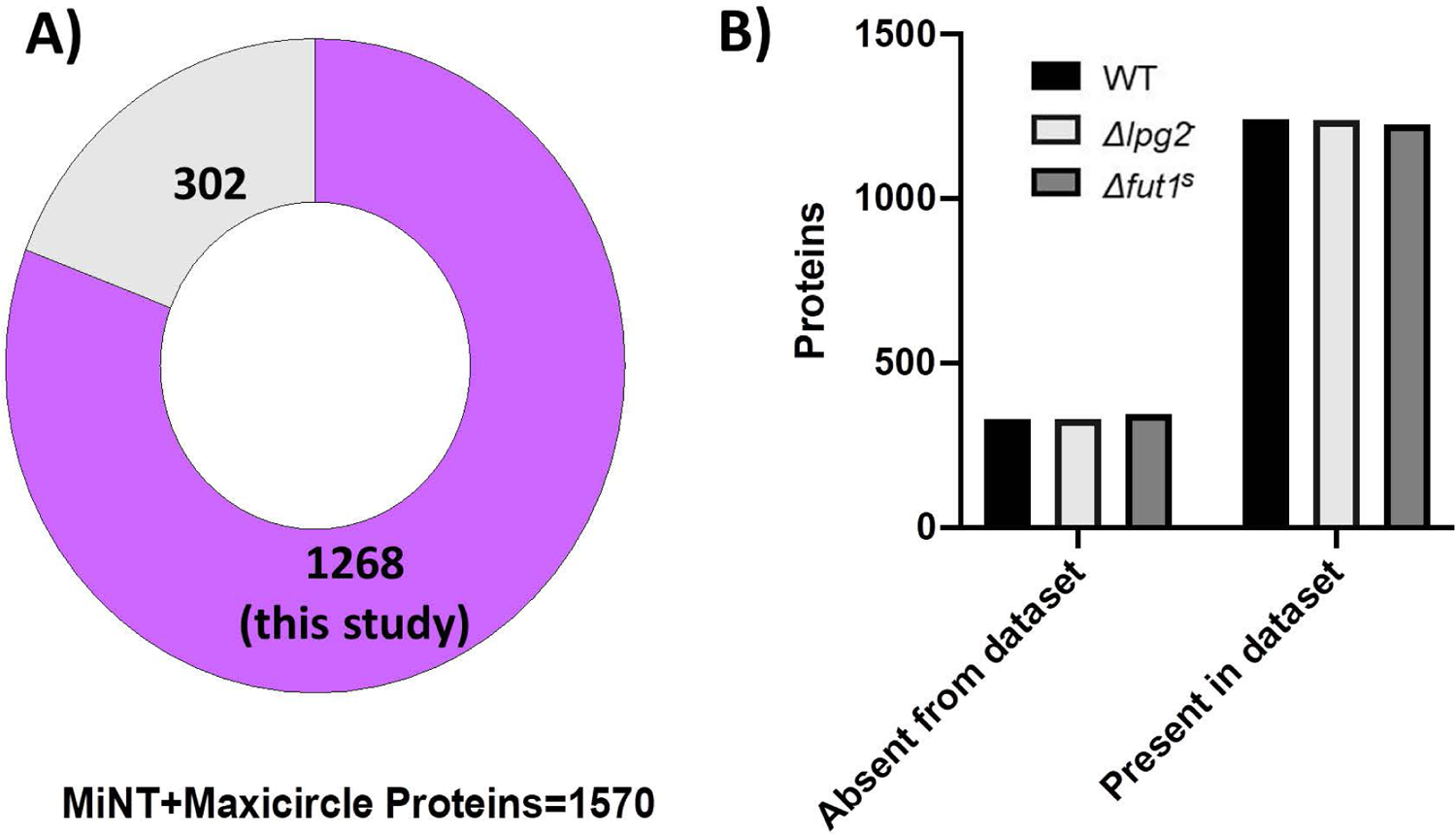
Representation <u>of the predicted *L. major* mitochondrial proteome.</u> Proteins from WT or mutant parasite lines (Table S3) were compared to proteins in predicted the nuclear encoded mitochondrial protein database MiNT, supplemented with maxicircle (mitochondrial) encoded proteins as described in the methods. A) 1268 proteins, representing 81%, of the mitochondrial proteome, were identified in this study. B) Distribution of mitochondrial proteins by cell line.

A gene ontology (GO) analysis was performed using PANTHER Classification System (Fig. S2, Table S4) (55). As seen previously (36), much of the *L. major* proteome consists of hypothetical proteins lacking annotation (over 60%; Fig. S2A), and GO analysis of annotated proteins showed considerably similarity amongst all three lines. For biological process (Fig. S2B), the categories with the highest number of genes were cellular process (GO:0009987), metabolic process (GO:0008152), localization (GO:0051179), and biological regulation (GO:0065007). For cellular component (Fig. S2C), the most commonly assigned categories were cellular anatomical entity (GO:0110165), intracellular (GO:0005622) and protein-containing complex (GO: 0032991). For molecular function (Fig. S2D) were catalytic activity (GO: 0003824) and binding (GO:0005488).

### Knockout of *FUT1* or *LPG2* has significant differential impact on *L. major* protein abundance

Quantitative differences amongst WT, *Δfut1**^s^*** or *Δlpg2^-^* parasite were assessed by ANOVA testing with an FDR of 0.05 applied to the protein abundance values (LFQ; Table S3), which were then normalized by Z-scoring to construct a heat map (Fig. 4). Clustering showed clear grouping of replicates within each line, as well as blocks encompassing 465 proteins differing significantly (Fig. 4). Of these, 107 increased and 142 decreased in *Δfut1^s^* only, 173 decreased and 13 increased in *Δlpg2^-^* only, and 30 which decreased in *Δfut1^s^* but increased in 2/4 of the *Δlpg2^-^* biological replicates (Tables S3, S5, S6; Fig. 4). Again, the majority of differentially expressed proteins were hypothetical proteins (254/465 total; Table S3).

**Figure 4.**
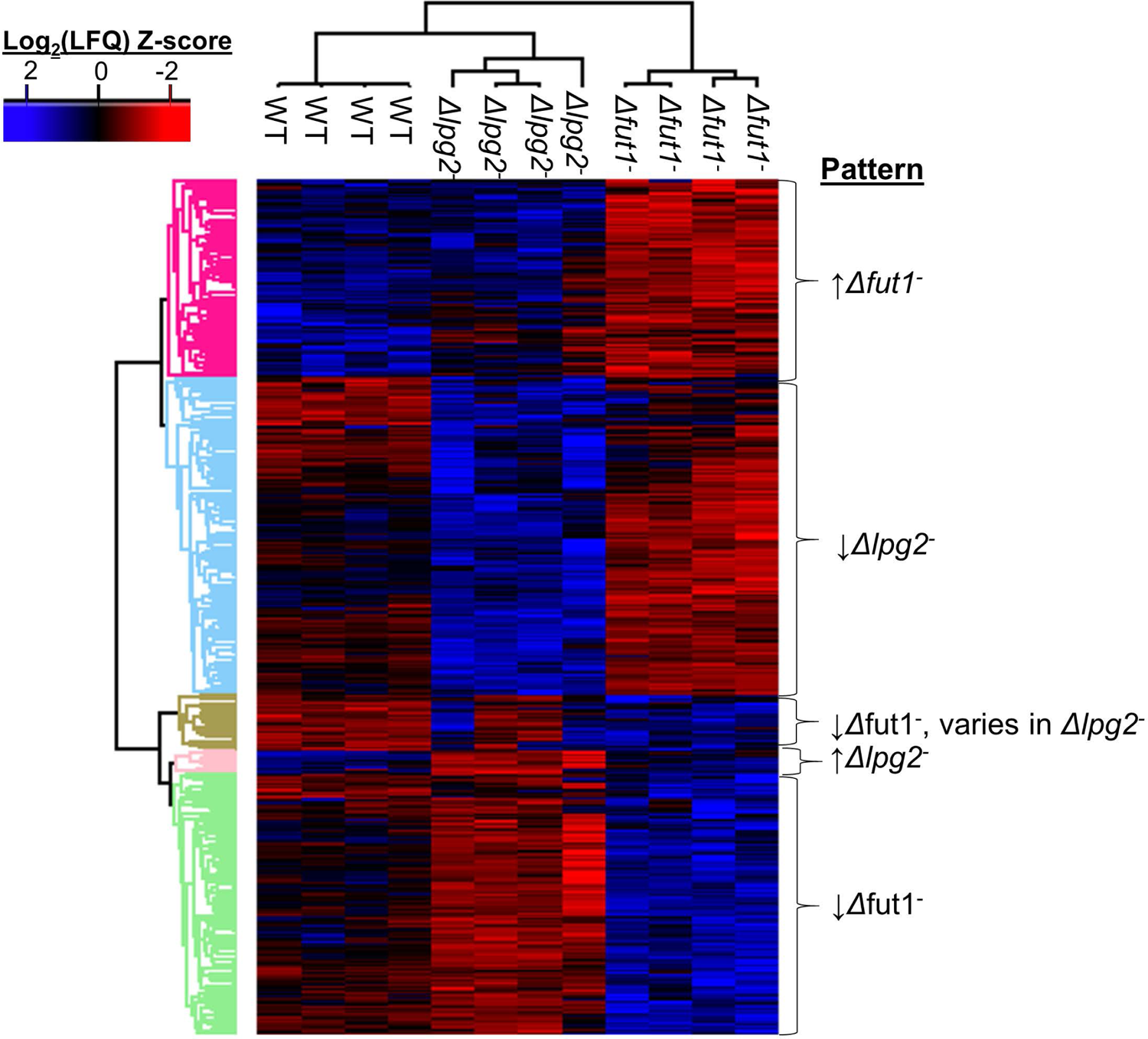
Patterns <u>of variation of significantly differentially expressed proteins amongst WT and *FUT1* and *LPG2* knockout *L. major*.</u> ANOVA testing using an FDR of 0.05 and an S0 of 1 was performed on LFQ values for the ‘high confidence’ proteome (Table S3), to identify proteins that are significantly different between WT, *Δfut1**^s^***, and *Δlpg2^-^*. From these data a heatmap was generated, clustering the parasite lines, each with four replicas (top dendrogram), and the differentially expressed genes (left dendrogram). The cluster properties are summarized on the right and the specific proteins included in each cluster can be found in Table S5 and S6.

SHERP, a stationary phase / metacyclic parasite marker (Sadlova et al. 2010), was not seen in any *Δlpg2*^-^ replicates, but was unexpectedly detected in log phase WT and *Δfut1^s^*parasites at similar levels (Table S1, S3). These results were corroborated by western blot analysis with anti-SHERP antisera (Fig. S3).

### Significantly different proteins accompanying the severe mitochondrial dysfunction of *Δfut1^s^*

The *Δfut1^s^* mutant shows considerably slower growth and numerous mitochondrial abnormalities; including lowered mitochondrial membrane potential, complete loss or loss in compactness of the kDNA network (30). From the cluster of 142 significantly less abundant proteins in *Δfut1**^s^***(Fig. 4; Table S5), 26 (18%) were predicted as mitochondrial, of which 13 were annotated (Table 2). Out of 107 proteins significantly more abundant in *Δfut1**^s^***, 15 were mitochondrial of which 2 were annotated (Table 2). The differentially expressed mitochondrial proteins are depicted in Fig. 5, many of which are positioned to contribute to *Δfut1**^s^*** mitochondrial dysfunction (Discussion).

**Figure 5.**
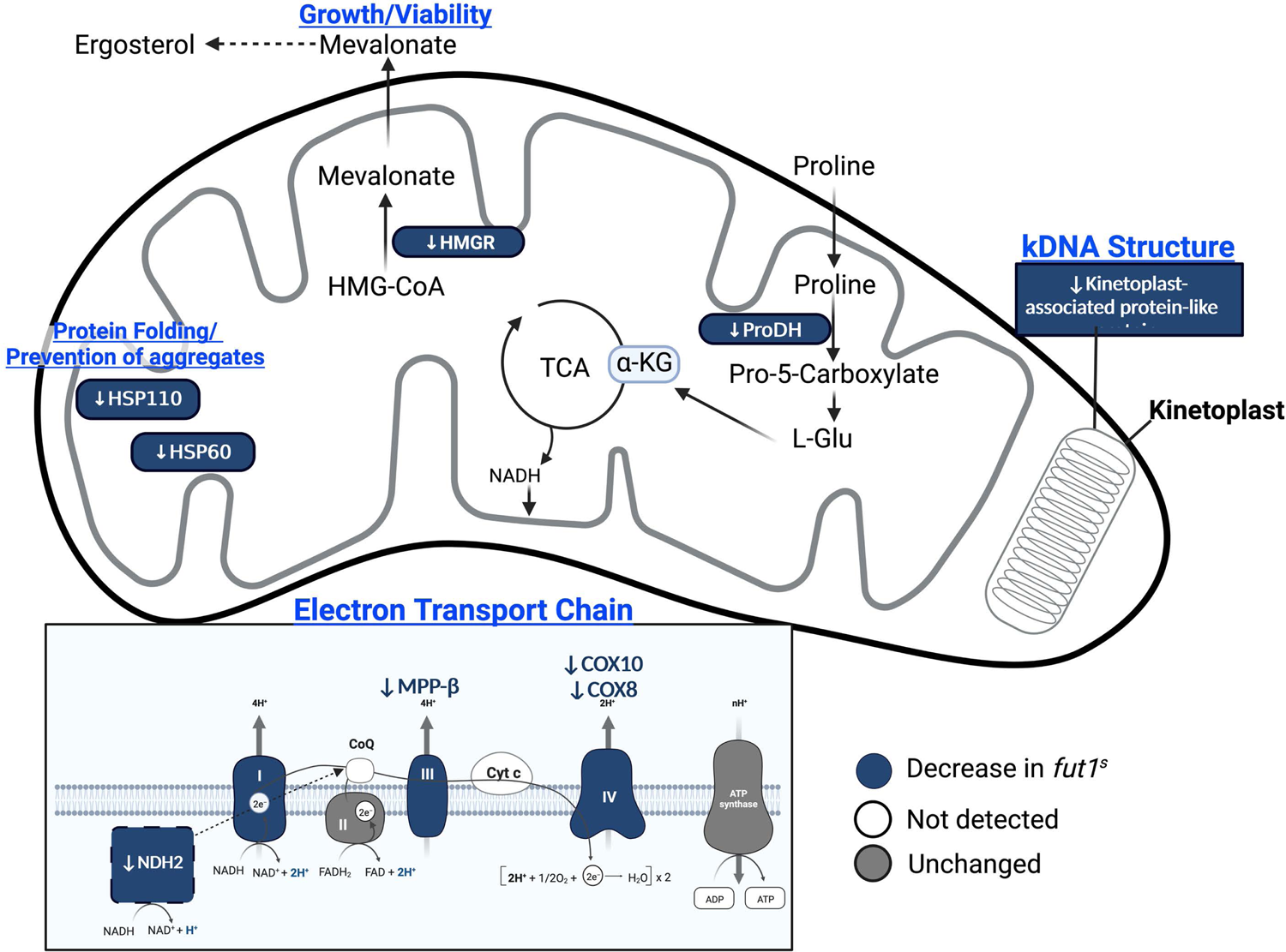
Schematic <u>overview of a *Leishmania* mitochondrion showing proteins with significantly altered</u> <u>expression in *Δfut1*</u>***^s^*** relative to WT *L. major*. The specific proteins are listed in Table 1. Only mitochondrial proteins showing differential expression (Table 2) are shown. In the inset depicting the electron transport chain, the proteins decreased in *Δfut1**^s^***are shown in blue, ones not detected in our dataset are shown in white, and ones that were unchanged are shown in gray. It should be noted that trypanosomatids possess a single mitochondrion per cell whose structure differs from the ‘typical metazoan’ mitochondrion shape depicted here.

**Table 2.** Mitochondrial Proteins Significantly changed in *Δfut1^s^.* Annotated mitochondrial proteins differing significantly in abundance by ANOVA tests between the *Δfut1**^s^*** vs. WT or *Δlpg2^-^* parasites are shown (Fig. 4, Tables S3, S5). Overall, there were 279 proteins in clusters where proteins decreased in *Δfut1^s^* (142 total; 26 mitochondrial of which 13 are unannotated), increased in *Δfut1^s^* (107 total; 17 mitochondrial of which 15 are unannotated), or decreased in *Δfut1**^s^***and varied in *Δlpg2-* (30 total; 13 mitochondrial of which 12 are unannotated).

| Cluster Pattern | Protein Name | TriTrypID | Difference log<sub>2</sub> (LFQ intensity) |  |  | Corrected ANOVA p-value |
| --- | --- | --- | --- | --- | --- | --- |
|  |  |  | *Δlpg2<sup>-</sup>* vs WT | *Δfut1<sup>s</sup>* vs WT | *Δfut1<sup>s</sup>* vs *Δlpg2<sup>-</sup>* |  |
| Down in *Δfut1<sup>s</sup>* | 3-hydroxy-3-methylglutaryl coenzyme A reductase (HMG-CoA | LmjF.30.3190 | 0.87 | -1.38 | -2.26 | 0.01 |
|  | Aldehyde dehydrogenase, mitochondrial | LmjF.25.1120 | 0.12 | -2.00 | -2.13 | 0.01 |
|  | Chaperonin HSP60, mitochondrial | LmjF.36.2030 | 0.80 | -0.81 | -1.61 | 0.04 |
|  | Kinetoplast-associated protein-like protein | LmjF.27.0240 | 1.60 | -3.80 | -5.40 | 0.00 |
|  | Metallo-peptidase, Clan ME, Family M16 | LmjF.35.1380 | 0.26 | -1.14 | -1.40 | 0.04 |
|  | Proline dehydrogenase (EC 1.5.5.2) | LmjF.26.1610 | 1.34 | -0.07 | -1.40 | 0.02 |
|  | Putative asparaginyl-tRNA synthetase | LmjF.34.2340 | 0.42 | -1.06 | -1.48 | 0.03 |
|  | Putative carbamoyl-phosphate synthase | LmjF.16.0590 | 0.56 | -0.69 | -1.25 | 0.05 |
|  | Putative cytochrome c oxidase VIII (COX VIII) (EC 1.9.3.1) | LmjF.31.1570 | 0.63 | -2.16 | -2.79 | 0.02 |
|  | *Putative cytochrome c oxidase subunit 10 (EC 1.9.3.1) | LmjF.23.0370 | -1.34 | -2.49 | -1.15 | 0.04 |
|  | Putative DNA-directed rna polymerase I largest subunit (EC 2.7.7.6) | LmjF.32.0650 | 0.72 | -0.69 | -1.42 | 0.04 |
|  | Putative heat shock protein | LmjF.18.1370 | 0.56 | -0.83 | -1.38 | 0.04 |
|  | Putative NADH dehydrogenase (EC 1.6.99.3) | LmjF.36.5380 | 0.47 | -1.20 | -1.67 | 0.04 |
|  | Putative translation initiation factor | LmjF.17.1290 | 0.78 | -0.59 | -1.36 | 0.05 |
| Up in *Δfut1<sup>s</sup>* | Inositol phosphosphingolipid phospholipase C-Like | LmjF.08.0200 | 0.09 | 2.00 | 1.91 | 0.01 |
|  | Putative 3,2-trans-enoyl-CoA isomerase mitochondrial (EC 1.1.1.35) | LmjF.31.2330 | -0.79 | 2.14 | 2.93 | 0.04 |

### Protein changes seen in the secretory pathway mutant *Δlpg2^-^*

Overall, there were 174 proteins significantly decreased in *Δlpg2^-^*, of which 89 are of unknown function, while 13 increased in *Δlpg2^-^* of which 6 are of unknown function (Fig. 4, Tables S3, S6). The predominance of ‘down regulated’ proteins in *Δlpg2^-^* relative to *Δfut1**^s^*** could be related to the relative health of the *Δlpg2^-^* mutant (this work and (19)), as many of the changes seen in *Δfut1**^s^*** involved stress responses arising from mitochondrial dysfunction. Notably the proteins altered in *Δlpg2^-^* mapped to a variety of cellular compartments and metabolic pathways (Tables S4, S6), suggesting that despite the normal growth of this mutant in culture, its impact was nonetheless profound. The broad impact of *LPG2*-dependent effects thus hinders efforts to pinpoint key ones most important to the biological alterations of most interest, such as “persistence without pathology”.

*LPG2* loss affected several proteins involved in the glycosylation pathways. Two examples are decreases of phosphomannomutase (PMM) and phosphoacetylglucosamine mutase (PAGM) which catalyze the reversible transfer of phosphate between C6 and C1 hydroxyl groups of mannose and N-acetylglucosamine (56). PMM in particular is necessary for establishing infection in macrophages (27).

### Potential interactions between *LPG2-* and *FUT1*-dependent pathways

While the proteins differentially altered in *Δlpg2*^-^ and *Δfut1^s^*were mostly quite distinct (Fig. 4; Tables S5, S6), in 2/4 replicate *Δlpg2*^-^ lines a small group of 30 proteins showed inverse regulation to changes observed in *Δfut1^s^*(Fig. 4; Table S5). Of these, 12 were unannotated predicted mitochondrial proteins. As the replicates were grown at the same time from the same inoculation under the same conditions, we cannot account for the variability. Changes in *Δfut1**^s^*** were also observed in proteins outside the mitochondrion (Table S5). Interestingly, these included increase of other glycosyltransferases including α-1,2-mannosyltransferase, phosphoglycan β 1,2 arabinosyltransferase, and phosphoglycan β 1,3 galactosyltransferase 3 (Table S5). We speculate that the cross-mutant effects arise from competition for GDP-fucose synthesized in the cytosol (28, 29), for transport and use by secretory pathway fucosyltransferases (dependent on LPG2) or mitochondrion (FUT1).

## Discussion

This work presents a high-coverage *L. major* proteomic dataset consisting of new mass spectrometry-based evidence for nearly 3,500 proteins beyond the 3,600 proteins typically identified experimentally in several *Leishmania* species including *L. major* by others (36). For quantitative analysis, the data were parsed to yield a high confidence data set of 5,841 proteins. Comparison showed the proteomes of WT, *Δlpg2^-^,* and *Δfut1^s^*parasites were highly congruent, but with significant quantitative variation (57). Over half of significantly changing proteins were uncharacterized hypothetical proteins.

### FUT1 and mitochondrial dysfunction

In addition to severely delayed growth, the *Δfut1^s^* mutant exhibits profound mitochondrial dysfunction, including loss of membrane potential, bloated cristae, presence of large aggregates, loss kDNA compactness and complete loss of the kDNA network in some parasites (30). Benefiting from the high coverage of mitochondrial proteins in our dataset (>80%; Fig. 3), we were able to survey the impact of *FUT1* deletion on these (Tables 2, S5; Fig 4). The *Δfut1^s^* mutant had significantly decreased levels of key of components of mitochondrial respiratory chain complexes III (MMP-β), and IV (cytochrome c oxidase; COX8 and COX10), (58) as well as the alternative NADH dehydrogenase (NDH2) (57, 59). *Δfut1^s^*alterations in several maxicircle (mitochondrial) encoded components of the respiratory chain could not be assessed as they were absent in our datasets. Collectively down-regulation of this pathway matches the changes in membrane potential and growth seen in *Δfut1^s^* (30).

*Δfut1^s^* showed down-regulation of kinetoplast-associated proteins (KAPs), histone-like proteins attributed to packaging mitochondrial DNA (kDNA) in trypanosomatids (60). Down-regulation of these has been associated with rearrangement of the kinetoplast structure, parasite growth defects, shrinkage and complete loss of the kDNA network (61, 62), all phenotypes seen in *Δfut1^s^* (30).

Enzymes impacted in *Δfut1^s^* include 3-hydroxy-3-methylglutaryl coenzyme A reductase (HMGR) and proline dehydrogenase (ProDH) (Table 2, S5; Fig. 5). HMGR is a key enzyme in the mevalonate pathway yielding isoprenoids important for viability and proliferation (63). ProDH mediates the first step in a pathway that leading to production of α-ketoglutarate entering the tricarboxylic acid cycle (64) and generates FADH_2_. which transfers electrons to coenzyme Q and the electron transport pathways (64). ProDH transcripts have been found to be present in higher levels in drug-resistant *Leishmania* (65).

Several proteins associated with stress responses were also impacted in *Δfut1^s^*, including mitochondrial heat shock proteins HSP60 and HSP110 (66). HSP60 acts as one of the principal mediators of protein homeostasis in the mitochondrial matrix, and in response to damaged or misfolded mitochondrial proteins after oxidative stress (66, 67). The inositol phosphosphingolipid phospholipase C-like (ISCL) protein was increased in *Δfut1^s^,* which contributes to parasite survival in acidic environments (68).

### A first-pass *L. major* glycoproteome, lacking detectable fucosylation

The FUT1 protein shows motifs characteristic of fucosyltransferases within the GT11 family and recombinant *Trypanosoma brucei* and *L. major* FUT1 were able to fucosylate various glycans *in vitro* (30, 31). Moreover, *Lmj* FUT1 was shown to fucosylate several peptides *in vitro* ((30); TbrFUT1 was not tested). The *in vivo* substrate of FUT1 has proven elusive in both parasite species, and the broad specificity of LmjFUT1 prompted us to search for protein O-fucosylation as part of this work. While glycopeptides were not detected with high confidence in the total proteomic studies, potentially reflecting technical considerations (69, 70), following HILIC enrichment we were able to identify 58 glycopeptides representing 49 *Leishmania* proteins, many of which were known or anticipated to be glycosylated, but many were novel (Table 1). Several other potential modifications were identified computationally, such as pyrophosphorylation (71), which will require experimental verification in the future. In *Trypanosoma cruzi,* HILIC enrichment similarly successfully identified a variety of N- and O-glycosylated proteins (72). Previous comprehensive studies of the *Leishmania* glycoproteome were lacking (50), mostly limited to a few glycoproteins such as GP63 (leishmanialysin). Despite the utility of the HILIC enrichment, many known glycosylated proteins of *Leishmania* were not detected, and thus the proteins in Table 1 should be regarded as a ‘first pass’ preliminary glycoproteome, confirmation of which will be needed in any event.

Notably, we did not find evidence for peptide O-fucose or fucosylated glycans. Although glycopeptide enrichment using HILIC can be utilized for O-glycoproteome characterization, it may bias for peptides with large glycans or multiple glycosylations (73). It is possible that any potential *FUT1*-dependent protein fucosylation may be in a form recalcitrant to the methods used, or that other parasite stages could yield different results. It is also possible that further ‘proteomic searches extending our ‘ultradeep’ effort might eventually yield proteins or modifications (including fucosylation) not identified here. Alternatively, while it is difficult to prove a negative, the possibility that the *in vivo* substrate of FUT1 may be glycans or glycolipids rather than proteins or glycoproteins seems at least as likely from the absence of detectable peptide fucosylation. Resolution of this question ultimately will depend on identification of the relevant FUT1-dependent glycoconjugate in the future.

### Summary and perspective

Proteins showing altered expression in *Δlpg2*^-^ could contribute to the ‘persistence without pathology’ phenotype seen in this mutant; however, as these parasites grow normally in culture we were challenged to compose a predictive query of the 186 differentially expressed candidates for those most likely to contribute to the phenotype. These data also raised the possibility that changes in the expression of one or more of these proteins could contribute to the recovery of amastigote virulence in *Δlpg2*^-^ revertants occasionally found in infected animals (23). Experimental tests of the role(s) of the *LPG2*-dependent proteome in the loss or recovery of amastigote virulence will be required to resolve this in the future. Similarly, *Δfut1^s^* parasites showed 249 differentially expressed proteins, including many known or potential mitochondrial proteins as expected from its strong impact on mitochondrial function, as well as a number of non-mitochondrial proteins. While our studies provoke many hypotheses for the roles of both FUT1 and LPG2 in parasite biology, genetic confirmation will be required in the future, as well as to establish whether the effects seen are direct or indirect consequences of gene ablation. Fortunately, with the advent of high-throughput knockout screening via CRISPR technology (74), probing the importance of this panoply of genes will be increasingly feasible.

## Methods

### Cell culture

All parasites were derivatives of *L. major* strain Fn (MHOM/IL/80/Fn). Parasites were grown as the promastigote form *in vitro* in complete medium 199 (M199) supplemented with 10% heat-inactivated FBS, 40mM HEPES pH 7.4, 100nM adenine, 1μg/mL biotin, 5μg/mL hemin, penicillin-streptomycin, and 2μg/mL biopterin. Parasites were seeded in 200 mL media at a density of 10^5^ /mL in roller bottles and allowed to grow until mid-log phase, corresponding to a density of 1-4 x 10^6^ / mL. Culture vessels were rotated at approximately 2 revolutions per minute using a cell production roller apparatus (Bellco Biotechnology). The *Δfut1^s^* segregant was described previously (30).

### Generation of *Δlpg2^-^* mutant line by CRISPR/Cas9 mutagenesis

Parasites were engineered to express a human codon-optimized *S.pyogenes* Cas9 and an LPG2 sgRNA (plasmid p63Phleo-HspCas9 or B7521; sgRNA synthesized from template B7617) expressed from a U6 promoter as described (75). To replace the *LPG2* open reading frame (ORF), homology directed repair (HDR) templates were designed containing a hygromycin B (*HYG*) resistance gene flanked by sequences matching the 30 nucleotides flanking the *LPG2* ORF. A total of 10 μg of HDR DNA were transfected into the *L. major* HSpCas9-B7617 parasites, which were then plated on semisolid M199 media containing 50 μg/ml hygromycin B. Numerous HYG^r^ colonies were obtained, of which 26 were tested and appeared to be WT/*Δlpg2*::*HYG* heterozygotes. Subsequently, parasites were transfected with a blastocidin (*BSD*) resistance gene HDR, bearing the 30 nt of *LPG2* flanking sequence as before. The HSpCas9-B7617-HYG^r^ parasites were transfected and plated onto semisolid media containing 50 μg/ml hygromycin B and 10 μg/ml blastocidin. A total of 22 colonies were tested and confirmed to be *Δlpg2::HYG* / *Δlpg2::BSD* knockouts by PCR tests (Fig. S1) and preliminary data showed them to be LPG-deficient as judged by their failure to agglutinate with monoclonal anti-PG antibody WIC79.3. One clone was selected for further study (SpCas9-B7617 B+H-c14.2), hereafter referred to as *Δlpg2^-^*.

### Generation of parasite lysates for proteomic analysis

Whole parasite lysates were prepared as described (76) with some modifications. After 3 washes with ice-cold PBS, parasites were resuspended in 1mL of ice-cold lysis buffer (6M guanidinium chloride, 100 mM Tris pH 8.5, 10 mM tris (2-carboxyethyl)phosphine (TCEP), 40mM 2-chloroacetamide (CAA), supplemented with a protease inhibitor tablet (Roche) and 0.2mg/mL 1,10 phenanthroline). Samples were immediately boiled at 95°C in a thermomixer at 1500 rpm. After cooling samples on ice for ten minutes, an additional boiling step was done at 95°C. One mg of protein was collected from four biological replicates. The samples were acetone precipitated with 4V ice-cold acetone at −20°C overnight, and then a second time under the same conditions for at least 4 hours. After centrifugation and supernatant removal, samples were air dried while being covered with a Kimwipe and stored at −80°C until ready for analysis.

### Digestion of *L. major* proteome samples

Precipitated protein pellets were resuspended in 50% trifluoroethanol and then heated at 50°C for 10 mins with shaking at 1000 rpm. Resuspended samples were then reduced / alkylated in a single step with 20mM tris(2-carboxyethyl)phosphine and 40mM chloroacetamide for 1hour in the dark. Samples were then diluted tenfold with 100 mM triethylamonium bicarbonate and digested with trypsin (1/100 w/w) overnight with shaking at 800 rpm. Digested samples were acidified to a final concentration of 0.5% formic acid and desalted with 50 mg C18 SEP-PAK columns (Waters Corporation, Milford, USA) according to the manufacturer instructions. Briefly, tC18 SEP-PAKs were conditioned with Buffer B (0.1% formic acid, FA, 80% acetonitrile, ACN), washed with 10 volumes of Buffer A* (0.1% trifluoroacetic acid, TFA, 2% ACN), sample loaded, column washed with 10 volumes of Buffer A* and bound peptides eluted with buffer B then dried down by vacuum centrifugation.

### High-pH Fractionation

Proteome samples were fractionated by basic reversed phase chromatography according to the protocol of Batth *et al* (77) with minor modifications. Briefly, peptides were resuspended in 1ml of Buffer A (5 mM ammonium formate, pH 10.5) and separated using a 1100 series HPLC (Agilent Technologies, CA) using a Gemini NX-C18 column (4.6 x 250 mm, 5 µm, Phenomenex, CA) at a flow rate of 1 ml/min. Separation was accomplished using a 90 min gradient with samples loaded on the column at 2% Buffer B (5 mM ammonium formate, 90% ACN, pH 10.5) for 3 mins. The concentration of Buffer B was then ramped from 2% to 28% B over 45 min, from 28% to 40% B over 5 min, from 40% to 80% B over 5 min. The gradient was held at 80% B for 2 mins and then the column regenerated by being returned to 2% B over 10 mins and held at 2% B for 10 min. Sixty one-minute fractions were collected. Every fifth fraction was combined to generate a total of 12 pooled fractions, which were concentrated by vacuum centrifugation, desalted using C_18_ stage tips and then subjected to mass spectrometric analysis.

### Proteomics analysis using reversed phase LC-MS

Pooled basic reverse phase fractions were re-suspended in Buffer A* (2% ACN, 0.1% TFA) and separated using a two-column chromatography set up composed of a PepMap100 C18 20 mm x 75 μm trap and a PepMap C18 500 mm x 75 μm analytical column (Thermo Fisher Scientific). Samples were concentrated onto the trap column at 5 μL/min for 5 minutes and infused into an Orbitrap Elite™ (Thermo Fisher Scientific). 125-minute gradients were run altering the buffer composition from 2% buffer B (80% ACN, 0.1% FA) to 28% B over 95 mins, then from 28% B to 40% B over 10 mins, then from 40% B to 80% B over 5 mins, the composition was held at 80% B for 3 mins, and then dropped to 2% B over 2 minutes and held at 2% B for another 10 mins. The Elite Orbitrap Mass Spectrometer was operated in a data-dependent mode automatically switching between the acquisition of a single Orbitrap MS scan (60k resolution) and a maximum of 10 MS-MS scans with each ion subjected to both a HCD (15k resolution NCE 40, maximum fill time 200 ms, AGC 5*10^4^) and ETD (Ion trap analyzed, ETD reaction time 100 ms with supplementary activation enabled, AGC 1*10^4^) scans.

### Proteomic analysis

Proteome samples were processed using MaxQuant (v1.6.3.4. (78)) and searched against the *Leishmania major* MHOM (Uniprot accession: UP000000542 containing 8038 proteins) and TriTrypDB *Leishmania major* Fn (TrypDB 37) databases. The reference proteome database was supplemented with predicted proteins from the kinetoplast maxicircle (mitochondrion) of *L. major* (LmjF00.0040, NADH dehydrogenase 7(MURF3,ND7); LmjF00.0050, Cytochrome oxidase 3; LmjF00.0060, CYb; LmjF00.0070, MURF4(A6); LmjF00.0080, MURF1; LmjF00.0100, NADH dehydrogenase subunit 1 (ND1); LmjF00.0110 Cytochrome oxidase 2 (CO2); LmjF00.0120, MURF2; LmjF00.0130, Cytochrome oxidase 1 (CO1); LmjF00.0150, NADH dehydrogenase 4 (ND4); LmjF00.0180, NADH dehydrogenase 5 (ND5)); due to extensive pan-RNA editing, other protein products that could not be reliably predicted were not be included. Searches were undertaken using “Trypsin” enzyme specificity with carbamidomethylation of cysteine as a fixed modification. Oxidation of methionine and dhex modifications of serine / threonine residues were included as variable modifications and a maximum of 2 missed cleavages allowed. To attempt to identify any potential complex fucosylation events dependent peptide searching was enabled. To ensure the inclusion of only high-quality peptide spectral matches (PSMs), a PSM false discovery rate (FDR) of 0.1% was set while an FDR of 1% was allowed at the protein level. To enhance the identification of peptides between samples, the Match between Runs option was enabled with a precursor match window set to 2 minutes and an alignment window of 20 minutes with the label free quantitation (LFQ) option enabled (79).

The result file was then uploaded to Perseus (V1.6.0.7) (82) for statistical analysis. Potential contaminants, proteins only identified by site, and reverse decoys were removed. For LFQ comparisons, values were log2(X) transformed and biological replicates were grouped. Data were parsed further by removing proteins not identified in at least two biological replicates in at least one cell line. Missing values were imputed based on the observed total peptide intensities with a range of 0.3σ and a downshift of 1.8σ using Perseus. A multiple sample (ANOVA) test with a permutation-based FDR set at 0.05 and s0 set at 1 was performed to identify significantly differentially abundant proteins between *Δfut1^s^*, WT, and *Δlpg2^-^*. A heat map of significantly different proteins was constructed after Z-score based normalization and Eucladian clustering of the transformed LFQ values (83).

### Glycopeptide enrichment by ZIC-HILIC

250 μg of digested and C18 SEP-PAKs cleaned up dried whole cell lysates were resuspended in 80% acetonitrile, 1% TFA and glycopeptides enriched using homemade ZIC-HILIC stage tips as previously described (48). Briefly, ZIC-HILIC columns were first conditioned with 80% acetonitrile, 1% TFA and then samples loaded onto columns before being washed with 80% acetonitrile, 1% TFA and glycopeptides eluted with Milli-Q water. Samples were dried and stored at −20C until undergoing LC-MS.

### LC-MS analysis of ZIC-HILIC enriched samples

ZIC-HILIC enriched samples were re-suspended in Buffer A* and separated using a two-column chromatography set up composed of a PepMap100 C18 20 mm x 75 μm trap and a PepMap C18 500 mm x 75 μm analytical column (Thermo Fisher Scientific) coupled to an Orbitrap Fusion™ Lumos™ Tribrid™ Mass Spectrometer (Thermo Fisher Scientific). ZIC-HILIC enriched samples were analyzed using 185-minute gradients. Separation gradients were run for each sample, altering the buffer composition from 2% Buffer B to 28% B over 106 or 166 minutes depending on the run length, then from 28% B to 40% B over 9 minutes, then from 40% B to 80% B over 3 minutes; the composition was held at 80% B for 2 minutes, and then dropped to 2% B over 2 minutes and held at 2% B for another 3 minutes. The Lumos™ Mass Spectrometer was operated in a data-dependent mode with a single Orbitrap MS scan (350-1800 m/z, maximal injection time of 50 ms, an AGC of maximum of 4*10^5^ ions and a resolution of 120k) acquired every 3 seconds followed by Orbitrap MS/MS HCD scans of precursors (NCE 30%, maximal injection time of 100 ms, an AGC set to a maximum of 1.0*10^5^ ions and a resolution of 15k). HCD scans containing the oxonium ions (204.0867; 138.0545 and 366.1396 m/z) triggered two additional product-dependent MS/MS scans of potential glycopeptides; an Orbitrap EThcD scan (NCE 15%, maximal injection time of 250 ms, AGC set to a maximum of 2*10^5^ ions with a resolution of 30k) and a ion trap CID scan (NCE 35%, maximal injection time of 40 ms, an AGC set to a maximum of 5*10^4^ ions). Data files were searched using MSFragger (v15, (46)) using the *Leishmania major* MHOM proteome (Uniprot accession: UP000000542). Open database searches were performed allowing modifications between −150 and 1000 Da on the deep proteome analysis with the “global.modsummary.tsv” used to assess the presence of fucosylated modifications. For the detection of glycoforms within ZIC-HILIC enrichments, open database searches were performed allowing modifications between −150 and 2000Da. The results from the ZIC-HILIC open searches were combined within R and only assignments with a MSFragger Expectation<0.001 and a delta mass > 140 Da.

## Data availability

All MS data, search results and R scripts have been deposited into the PRIDE ProteomeXchange Consortium repository (80, 81) with identifiers PXD015966 or PXD035738.

## Acknowledgements

The authors would like to acknowledge Igor Almeida for discussions, and Deborah Dobson and Suzanne Hickerson for sharing preliminary data on the LPG status and mouse infectivity of *L. major* Fn *Δlpg2^-^.* Funding was provided by R01 AI031078 as well as the NIAID R01 research supplement to promote diversity in health-related research (G.P.). This work was supported by National Health and Medical Research Council of Australia (NHMRC) project grants awarded to NES (APP1100164). We would like to thank the Melbourne Mass Spectrometry and Proteomics Facility of The Bio21 Molecular Science and Biotechnology Institute at The University of Melbourne for the support of mass spectrometry analysis. We thank Greg Matlashewski for providing plasmids used in CRISPR/Cas9 editing and the members of our group for discussions.

## Author contributions

GP – investigation, data analysis, writing manuscript. NES – investigation, data analysis, resources, funding. LFL – investigation. SMB – data analysis, writing, resources, funding.

## Abbreviations

LPG: lipophosphoglycan

LC-MS/MS: liquid chromatography tandem mass spectrometry

dhex: deoxyhexose.

## Supplementary information

### Supplementary Tables

Please note that due to their size, Tables S1-S4 are included as on-line supplementary information.

**Supplementary Table S1.**

<u>Total *L. major* proteome.</u> All proteins identified by one or more peptide in WT, *Δlpg2*^-^ or *Δfut1*^s^ total or HILIC-enriched datasets were included. A summary comparing this dataset with previous work is found in Fig. 2A. The heading terms are defined below

- **<u>Protein Names:</u>** Names of proteins contained within group based on shared identified peptides
- **<u>Gene names:</u>** Names of genes associated with identified proteins
- **<u>Protein IDs:</u>** All proteins consistent with identification criteria. Matches were made to a database composed of TriTrypDB and UniProt identifiers.
- **<u>Majority Protein IDs:</u>** Protein(s) matching at least half the peptides matched to the protein group. These IDs were used for analyses and data interpretation.
- **<u>TriTryp IDs</u>**: Identifiers from TriTrypDB database associated with majority protein IDs
- **<u>No Prior MS evidence listed:</u>** Protein groups identified in this study, that had no MS-based evidence listed on TriTrypDB for *L. major*
- **<u>Prior MS evidence listed:</u>** Protein groups identified in this study, that had MS-based evidence listed on TriTrypDB for *L. major*
- **<u>Retained for analysis of biological significance:</u>** Proteins used for downstream analyses to determine biological significance of *LPG2* or *FUT1* deletion. Only proteins that were detected in at least two biological replicates within one or more parasite line were considered.
- **<u>Log2 (LFQ intensity):</u>** relative label-free quantitative value across all samples (LFQ intensity) after log2 transformation
- **<u>Peptides:</u>** Number of peptides associated with the protein(s)
- **<u>Razor + Unique peptides:</u>** Razor peptides are found in more than one protein group but assigned to the group with the highest number of identified peptides.
- **<u>Unique peptides:</u>** Peptides matching only one protein sequence within a group
- **<u>Sequence coverage [%]:</u>** Percent sequence coverage by identified peptides belonging to the best protein sequence in the group
- **<u>Unique + razor sequence coverage [%]:</u>** Coverage of protein sequence based on both unique and razor peptides identified belonging to the best protein sequence in the group
- **<u>Mol. Weight [kDa]:</u>** Molecular weight corresponding to the best protein in the group
- **<u>Q-value:</u>** False discovery rate within the protein group
- **<u>Score:</u>** Andromeda score measuring efficiency of matching theoretical fragment masses to the acquired spectra. The value is calculated as the -log_10_ (probability match is acquired by chance).
- **<u>Intensity:</u>** Sum of intensities for all peptides corresponding to the protein group
- **<u>MS/MS count:</u>** Number of spectra for the protein group

**Supplementary Table S2.** <u>Peptides and proteins identified by MS following HILIC enrichment</u>. The first tab shows peptides, the second tab shows modified peptides collapsed by modification and/or protein; the third tab shows N-linked glycopeptides, the fourth tab shows N-linked glycoproteins, and the fifth tab shows pyrophosphorylation or unidentified modifications. A summary of those glycoproteins showing database annotations is found in Table 1.

**Supplementary Table S3.** <u>High confidence proteome of *L. majo*</u>*r*. The total experimental proteome (Table S1) was parsed to retain only those proteins present in 2/4 replicas in one or more lines. Their representation amongst lines appears in Fig. 2B.

**Supplementary Table S4.** <u>Gene ontology assignments</u>. A summary of these appears in Fig S2.

### Supplementary Figures

**Supplementary Figure 1.**
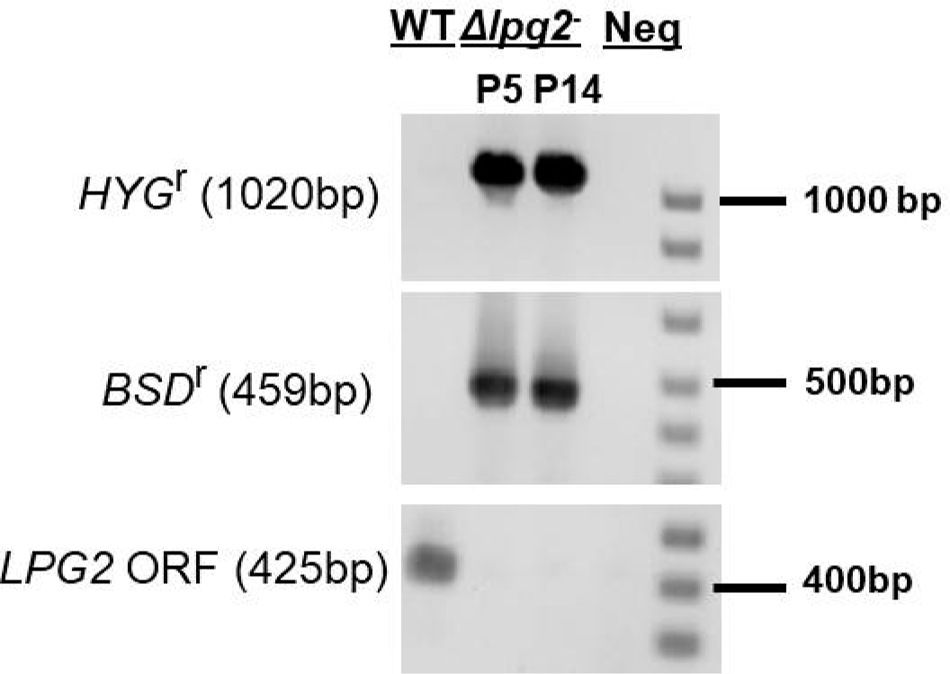
PCR confirmation of *L.major* Fn homozygous *LPG2* knockouts (*Δlpg2^-^)* obtained by <u>CRISPR/Cas9 mutagenesis</u>. Template DNAs used were WT and two different *Δlpg2^-^* clonal lines; ‘Neg’ corresponds to no template DNA added. Clone P14 was taken for proteomics studies presented here. The upper panel show amplification with the hygromycin B drug resistance marker (HYG^r^) with primers SMB2891 (5’-GGAGGACCCGGGCCACCATGAAAAAGCCTGAACTCACCG and SMB2892 (5’-GAGGATCTAGACTATTCCTTTGCCCTCGGACGA. The middle panel shows amplification with the blastocidin resistance marker (BSD^r^) with primers SMB7919 (5’-CCAACCGAAAGAATTGCATCAGCAACTGTC CCACCATGGCCAAGCCTTTGTC) and SMB7920 (5’-CCCTTCTACGACTGCGGCTAACAACGGTGATTAGCCCTCCCACACATAACCA). The lower panel shows amplification for the *LPG2* ORF with primers SMB7914 (5’-TCTGTCAGTAACTCGATCGGCC) and SMB7915 (5’-CGTCTTGCCGGTCTGCTGCATC).

**Supplementary Figure 2.**
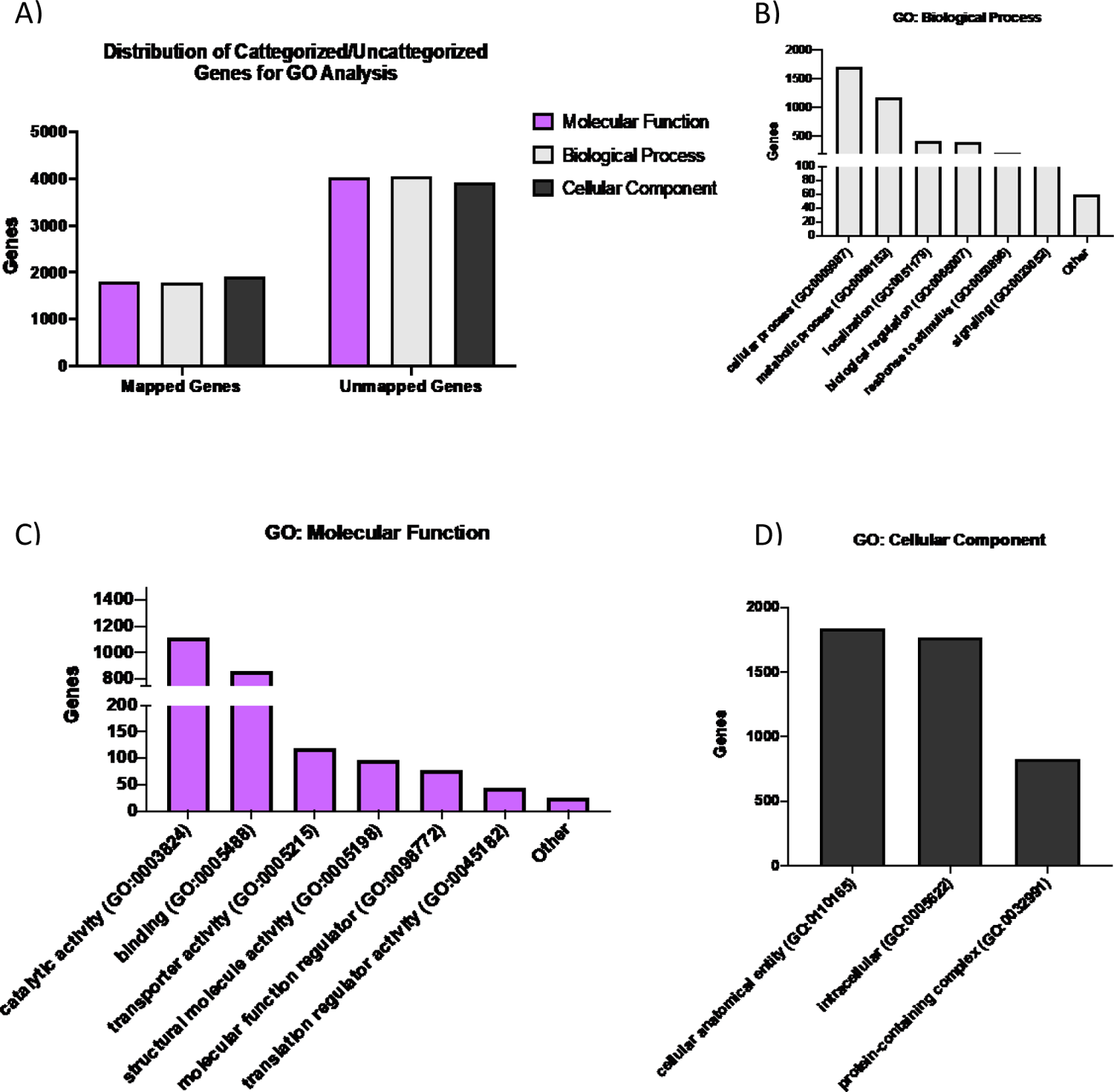
<u>Gene ontology (GO) analysis of the experimental *L. major* proteome</u>. The gene ontology assignments were determined using the PANTHER classification system. A) Numbers of proteins able to be assigned or not. Classification of proteomic datasets from each cell line by B) biological process, C) molecular function, and D) cellular component.

**Supplementary Figure 3.**
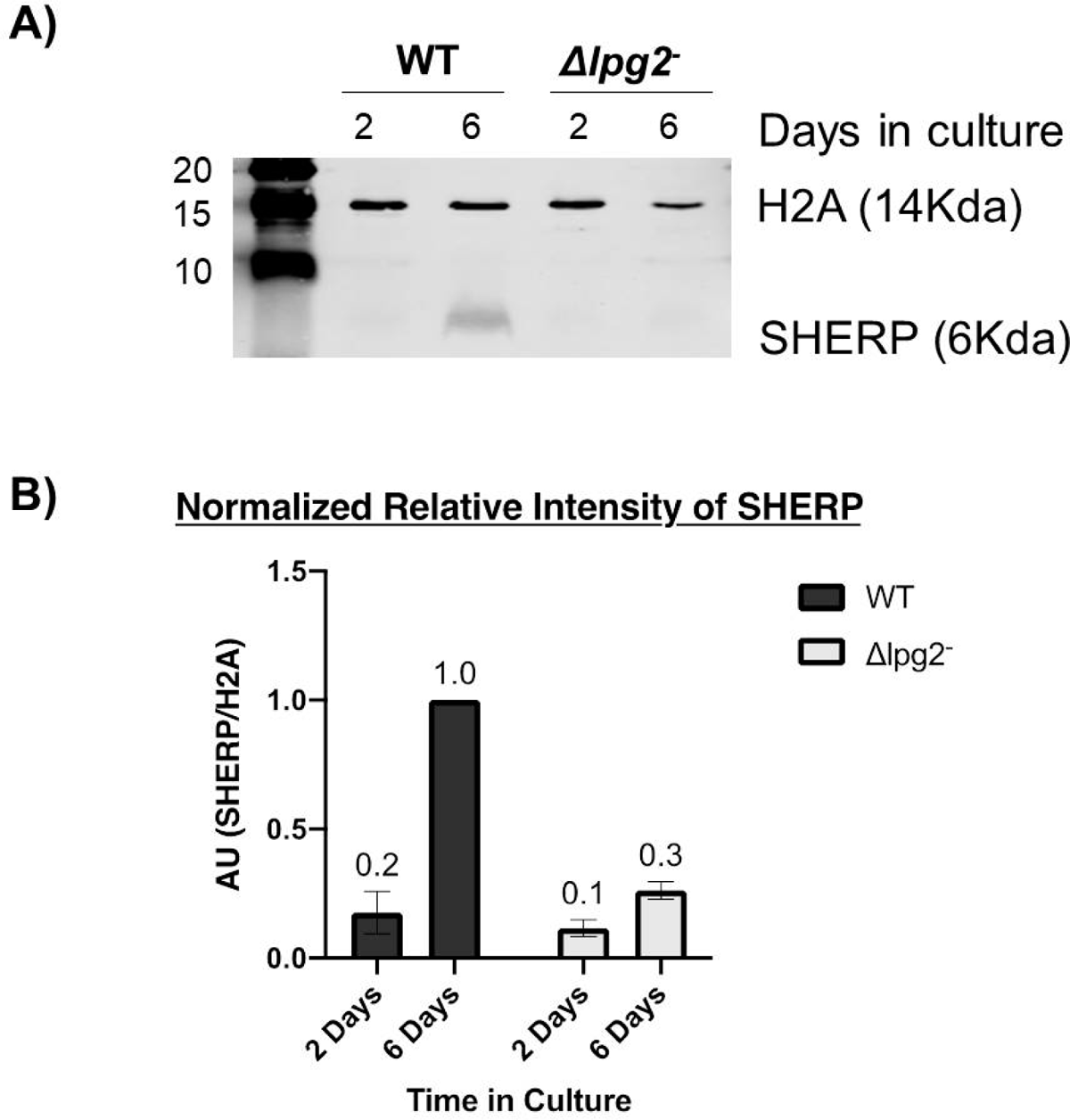
<u>Western blot analysis of SHERP expression in WT and *Δlpg2-L. major*</u> A) Western blot of 2 and 6 day old culture of WT and Δlpg2-parasites probing for SHERP and H2A loading control. B) Densitometry results from two biological replicates.

**Table S5:**
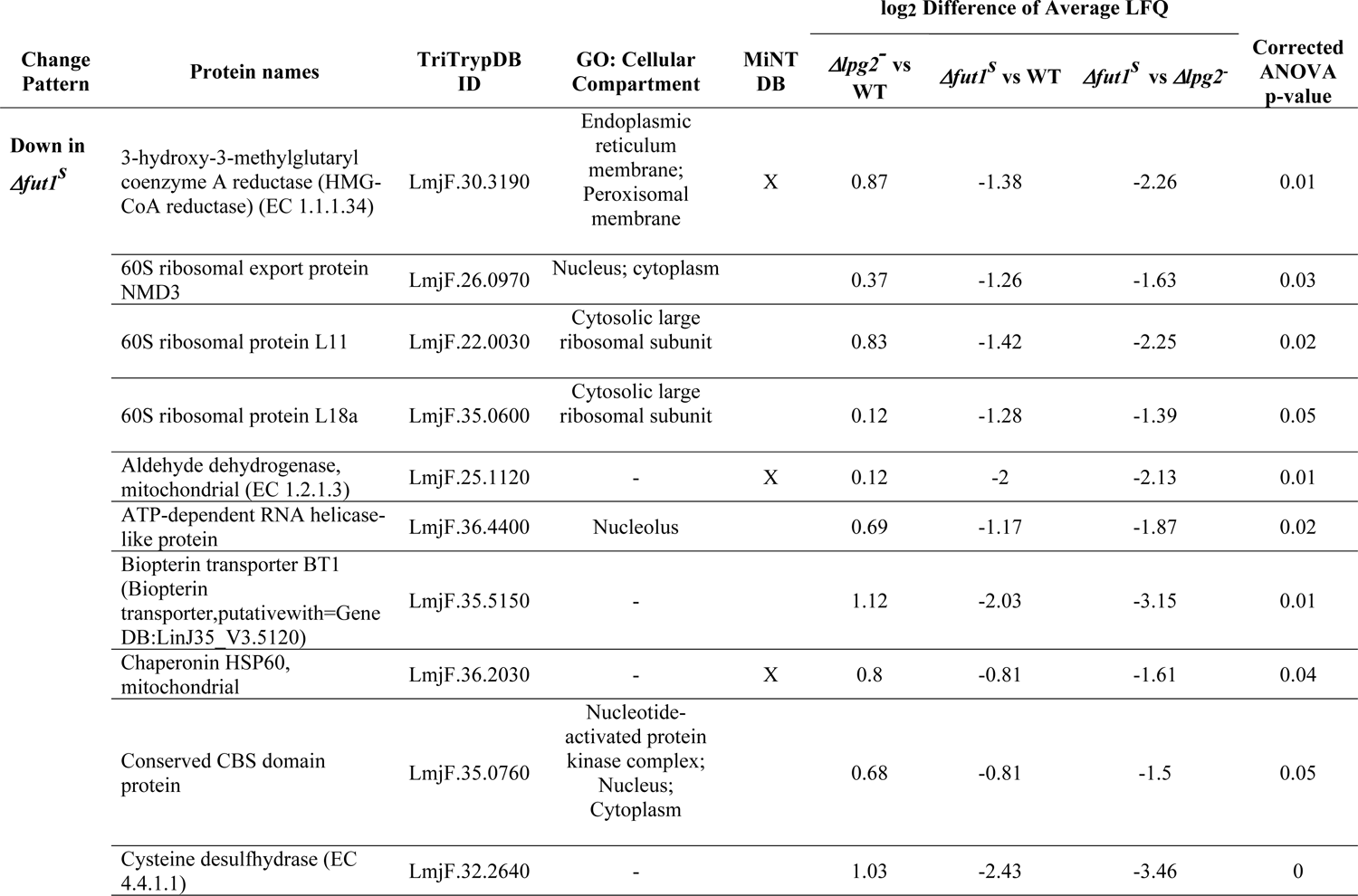
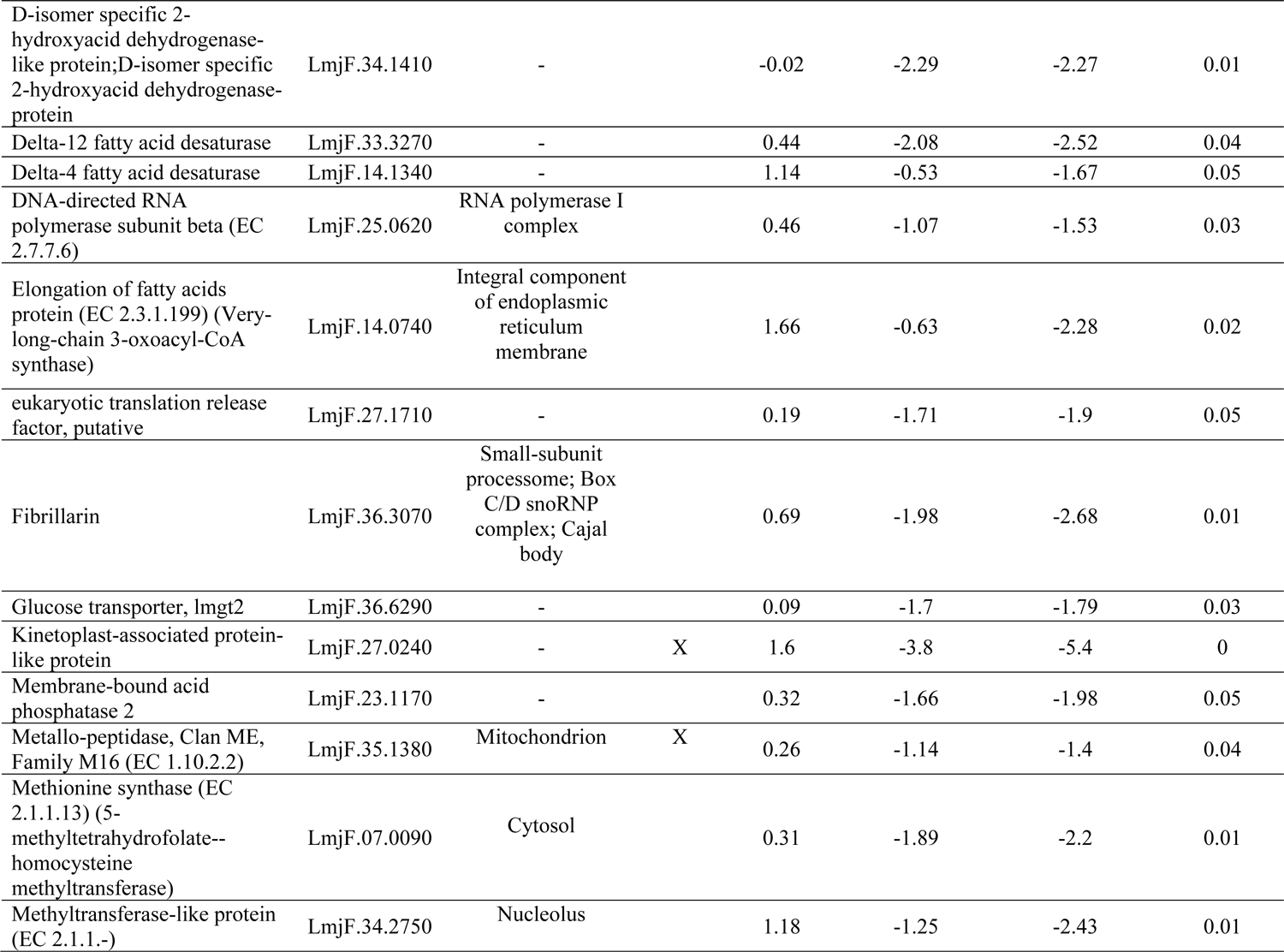
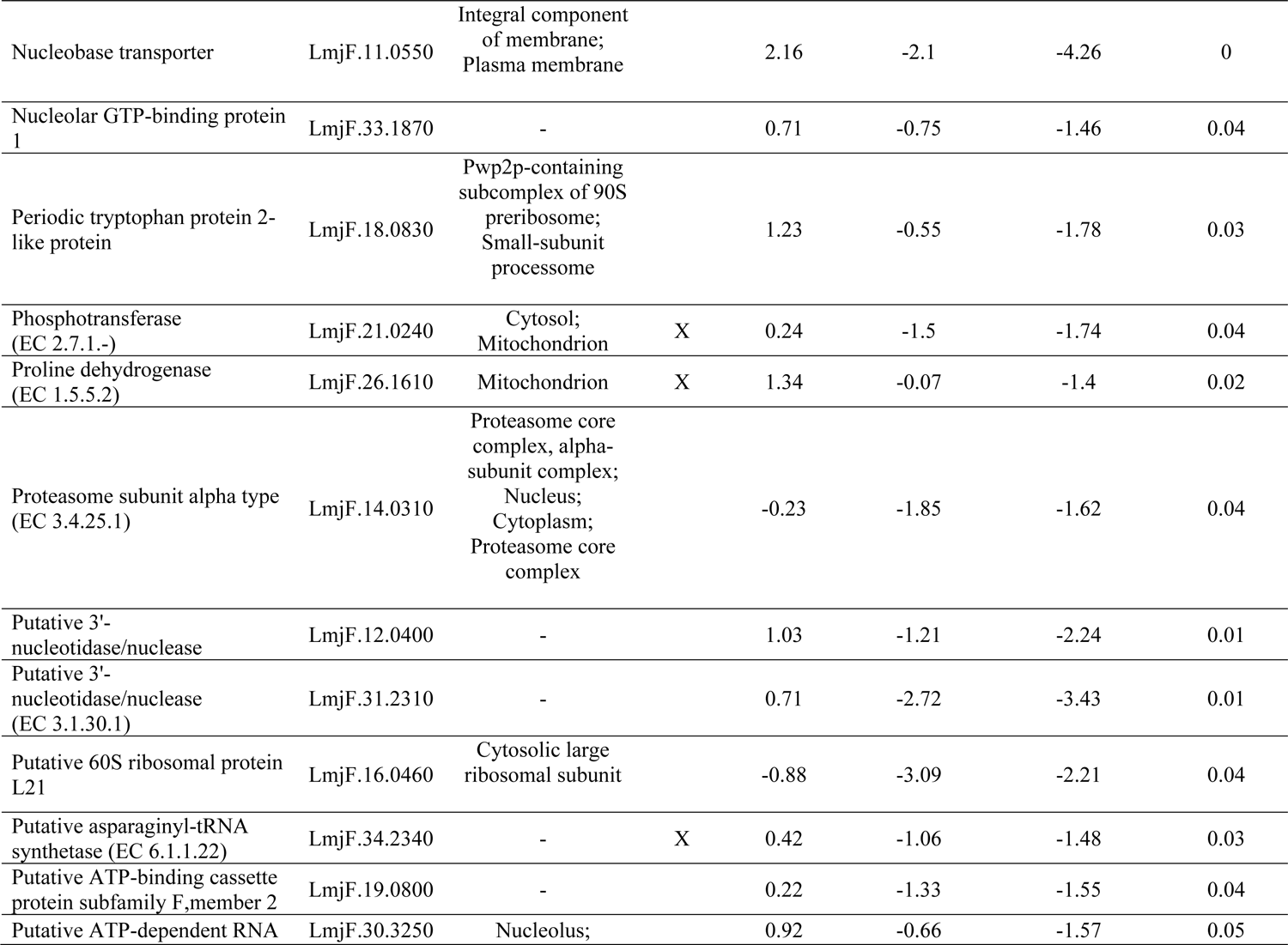
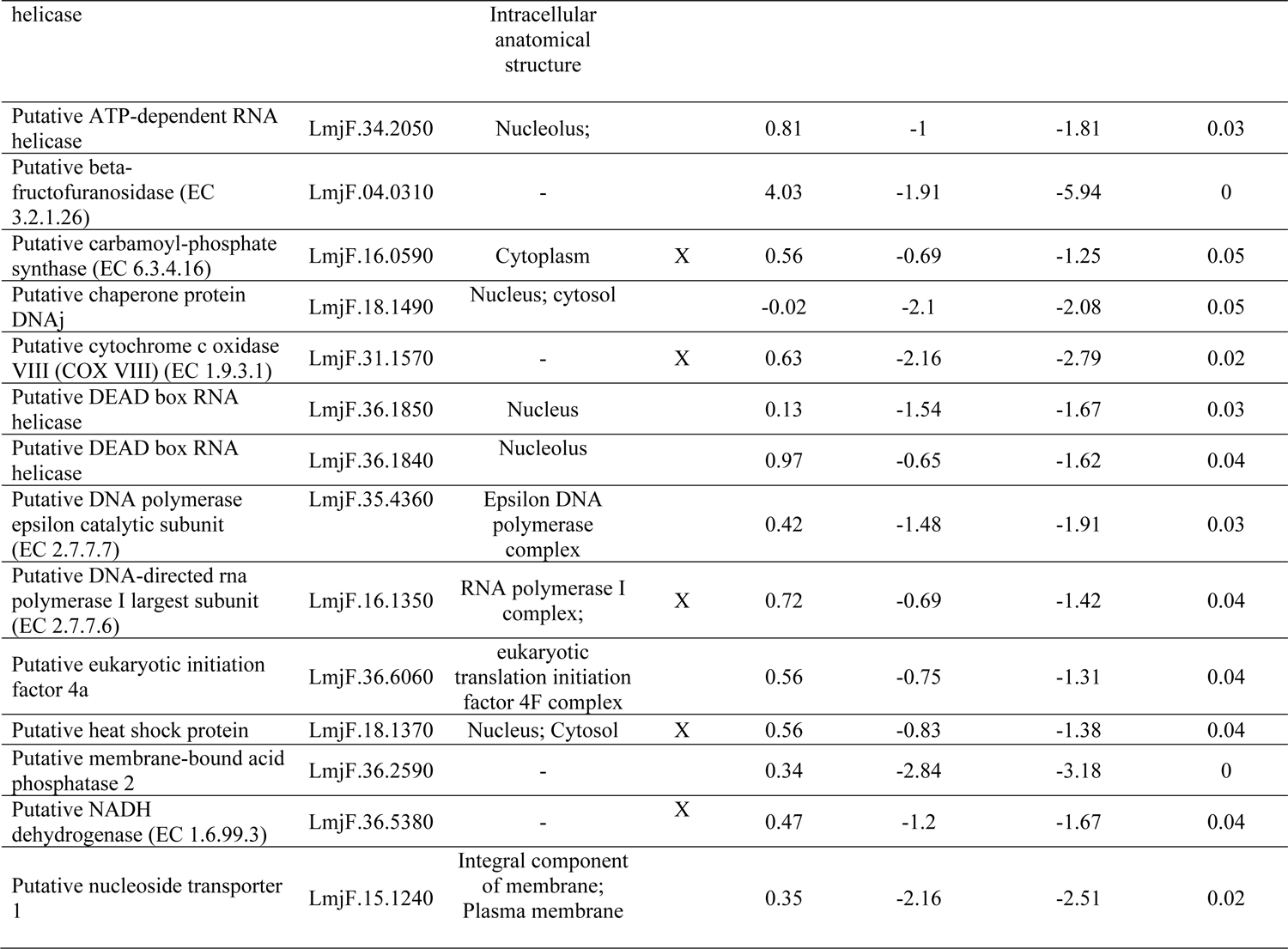
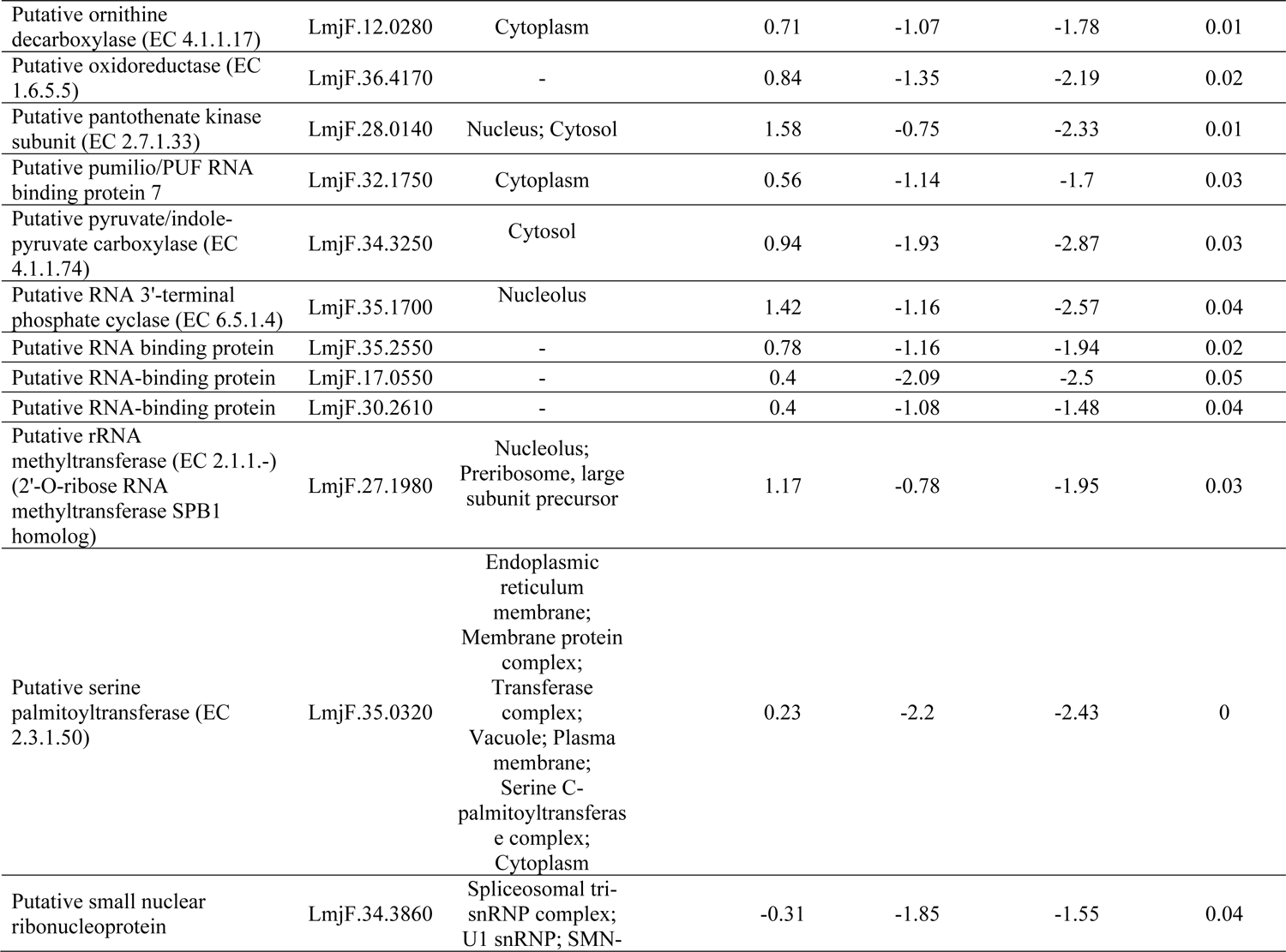
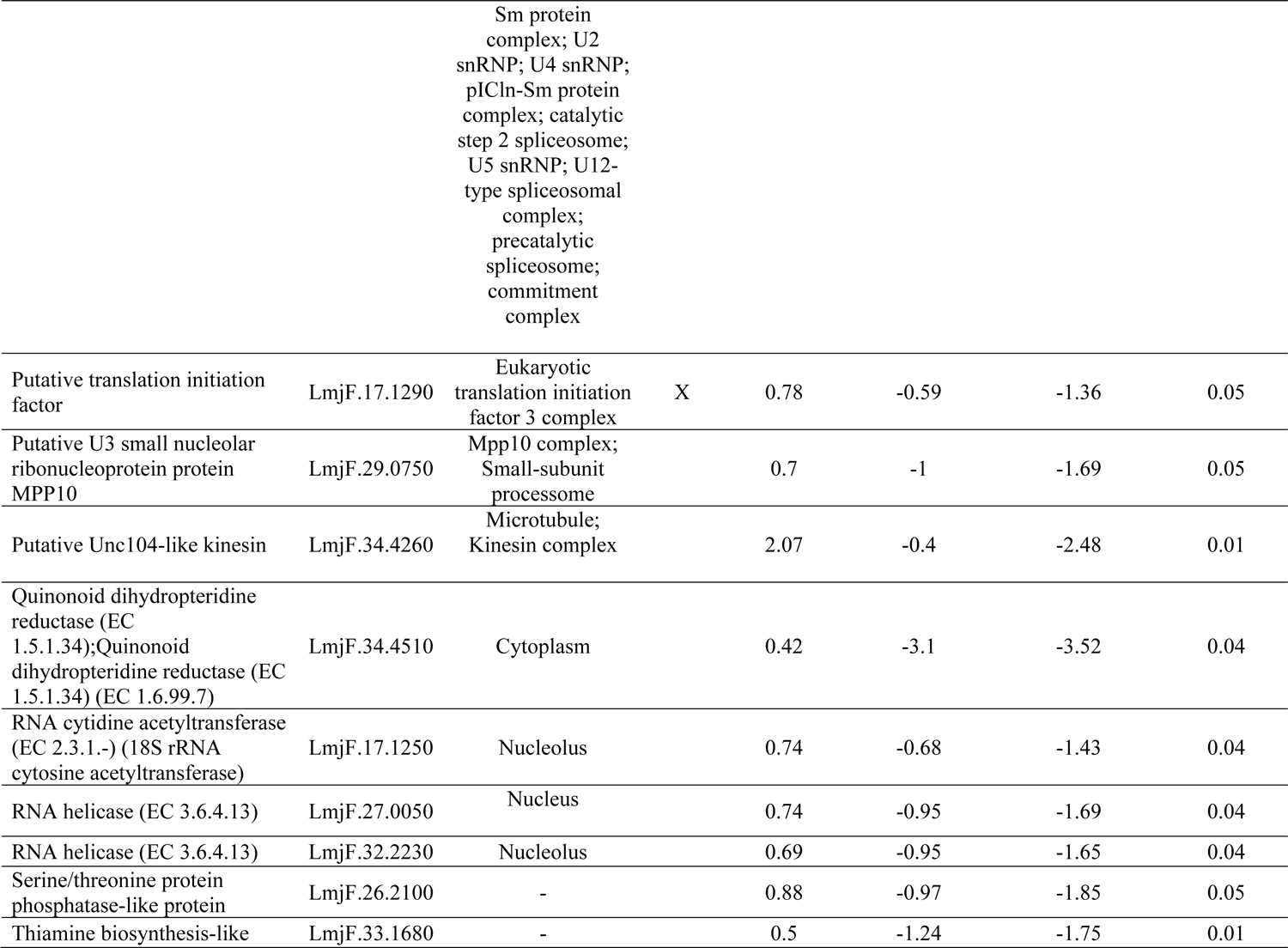
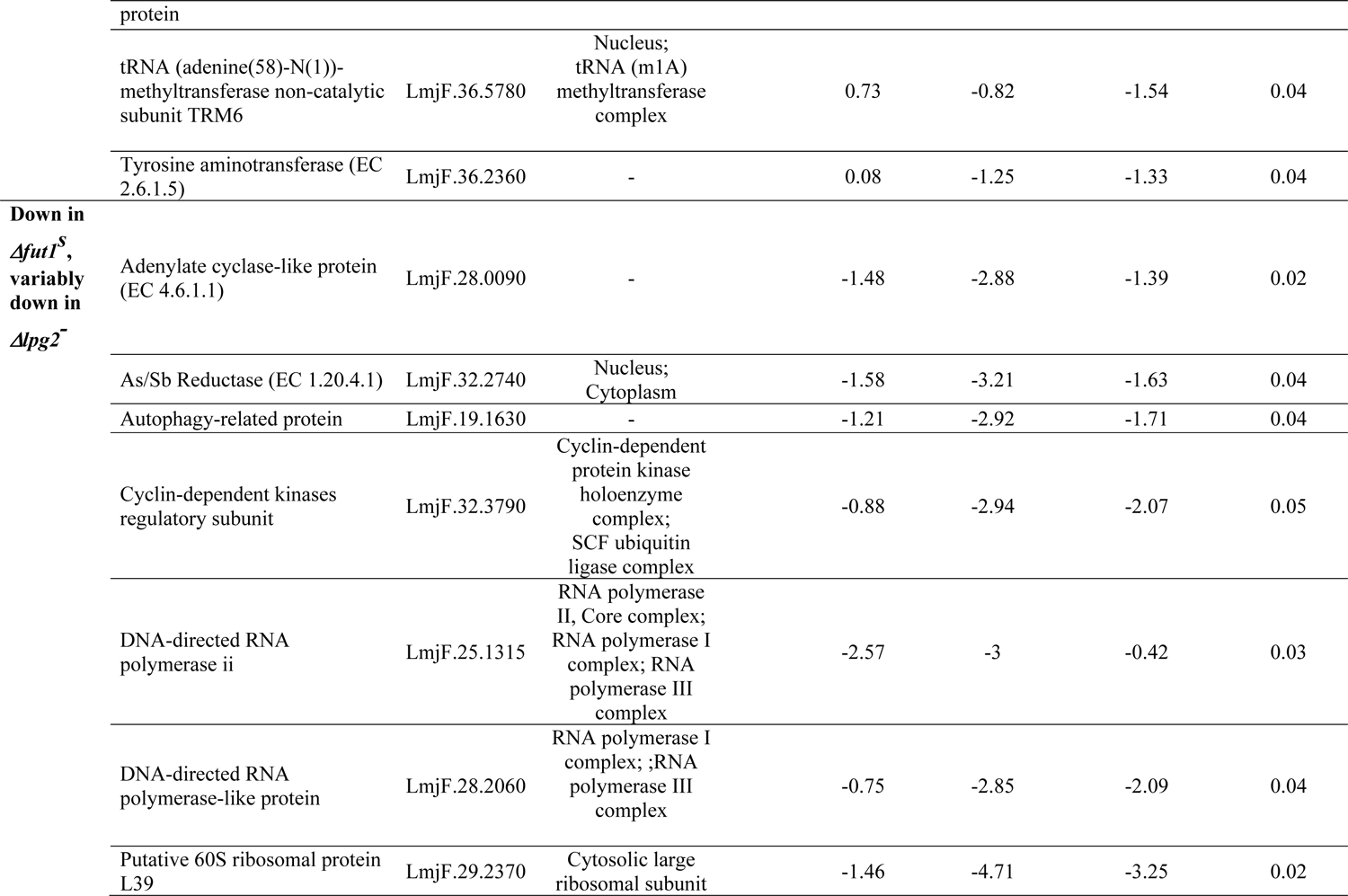
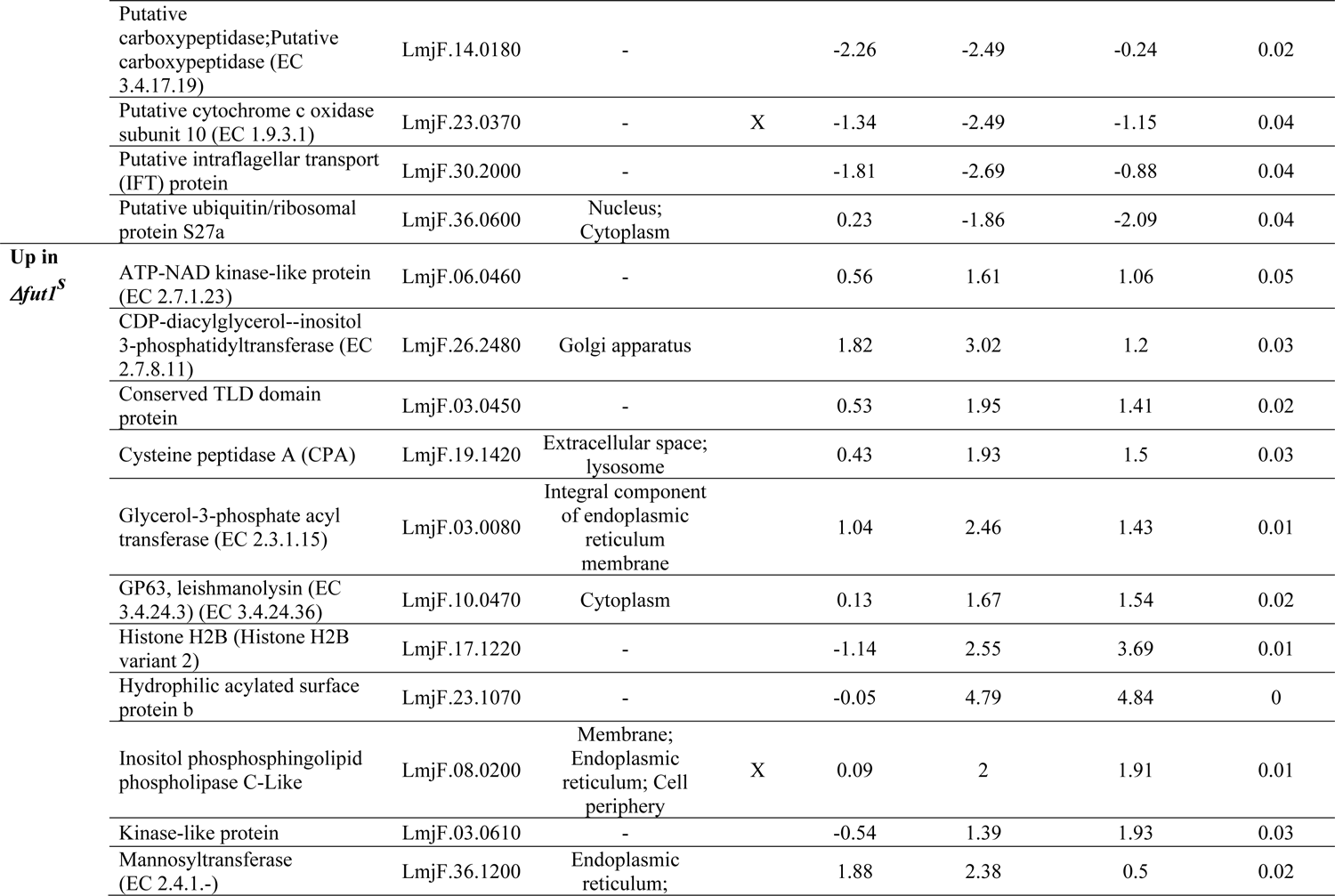
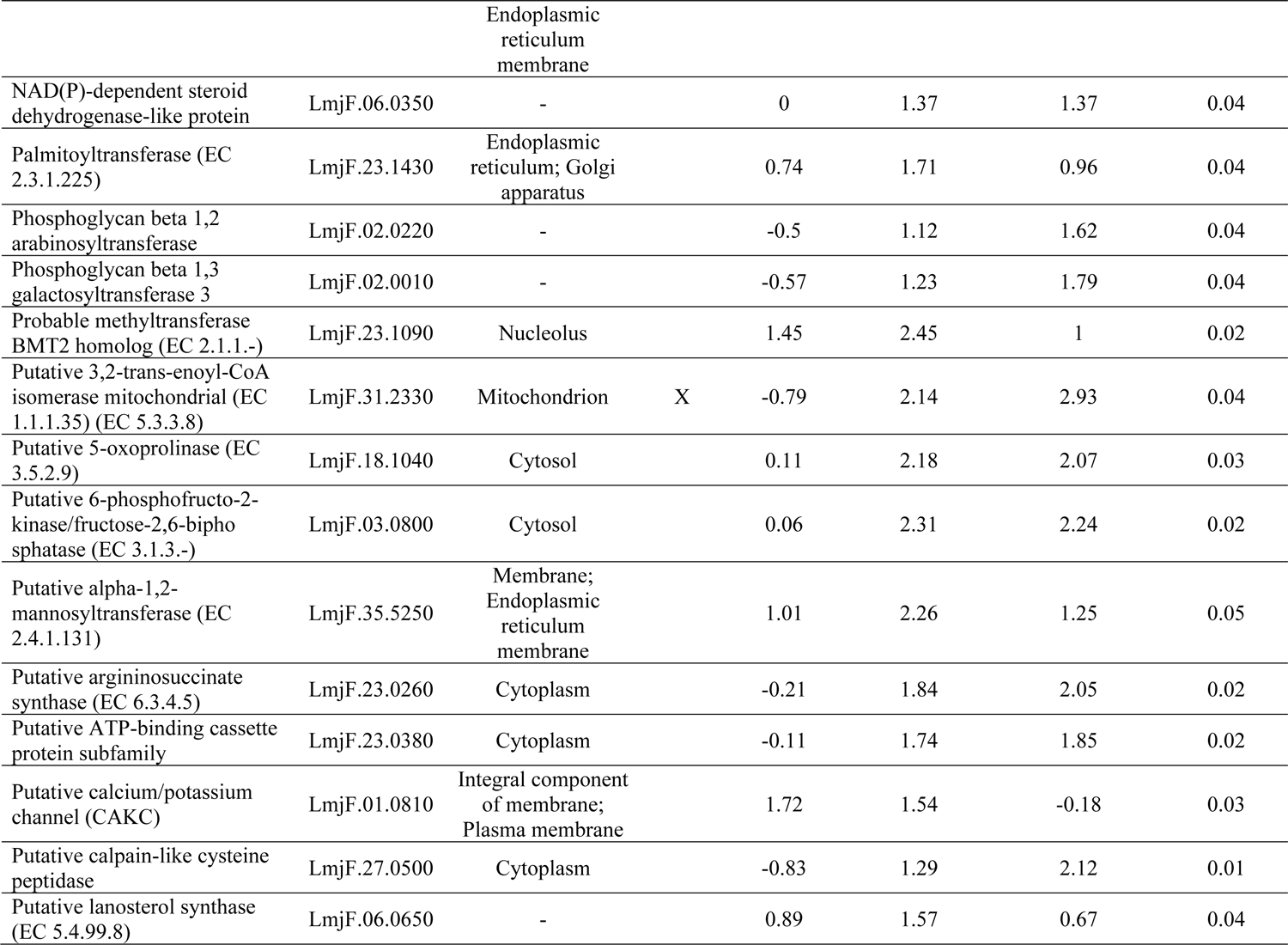
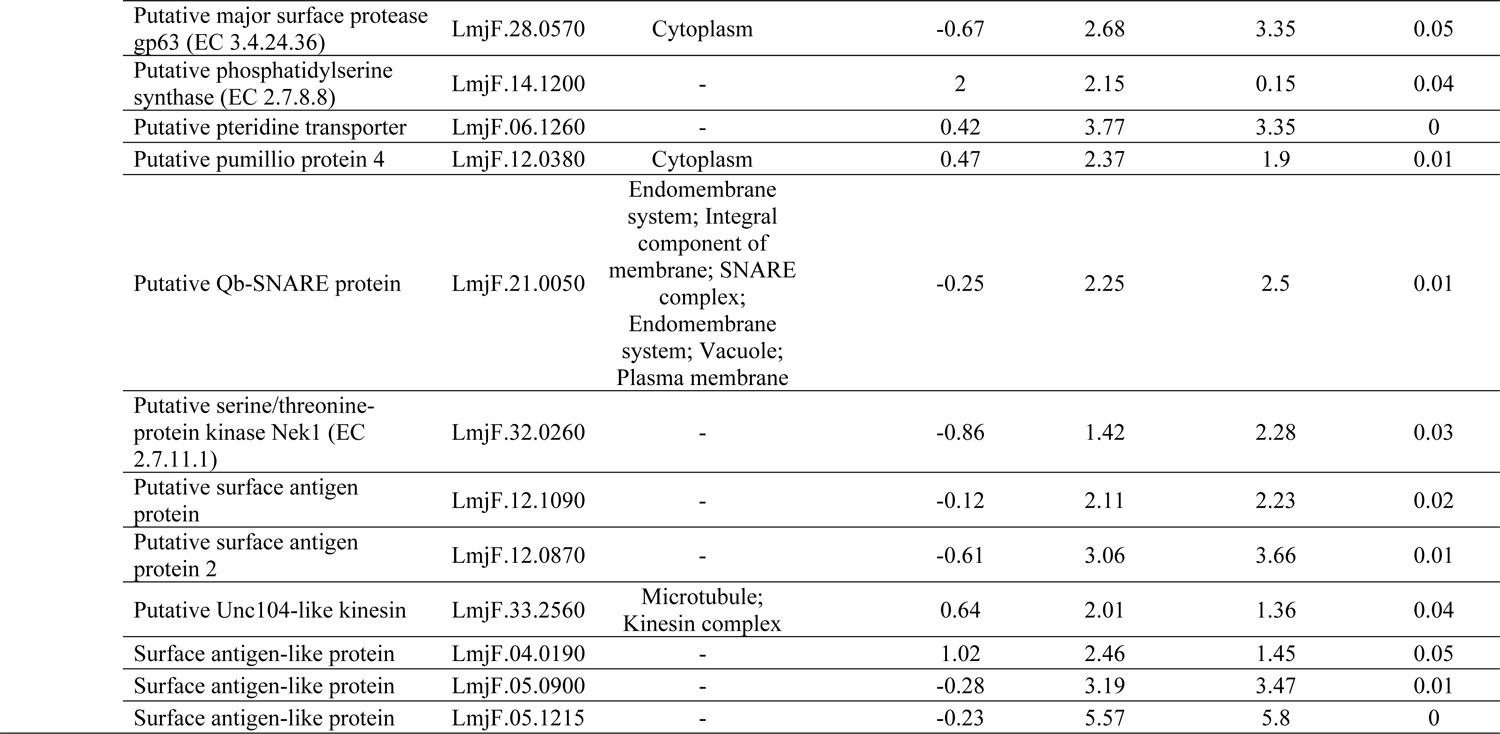
Annotated Proteins Significantly Affected in *Δfut1^s^*. Proteins determined to differ significantly in abundance between the *Δfut1**^s^**, Δlpg2^-^,* and WT parasite lines were evaluated by ANOVA; those proteins within clusters changing significantly in *Δfut1^s^*are shown here. Overall, there were 279 proteins in clusters where proteins decreased in *Δfut1^s^* (142 total of which 70 are of unknown function), increased in *Δfut1^s^* (107 total of which 70 are of unknown function), or decreased in *Δfut1**^s^*** and varieable impacted in *Δlpg2-* (30 total of which 19 are of unknown function). Proteins present in the MiNT database are indicated, as are the correspoding cellular component GO terms when available.

**Table S6:**
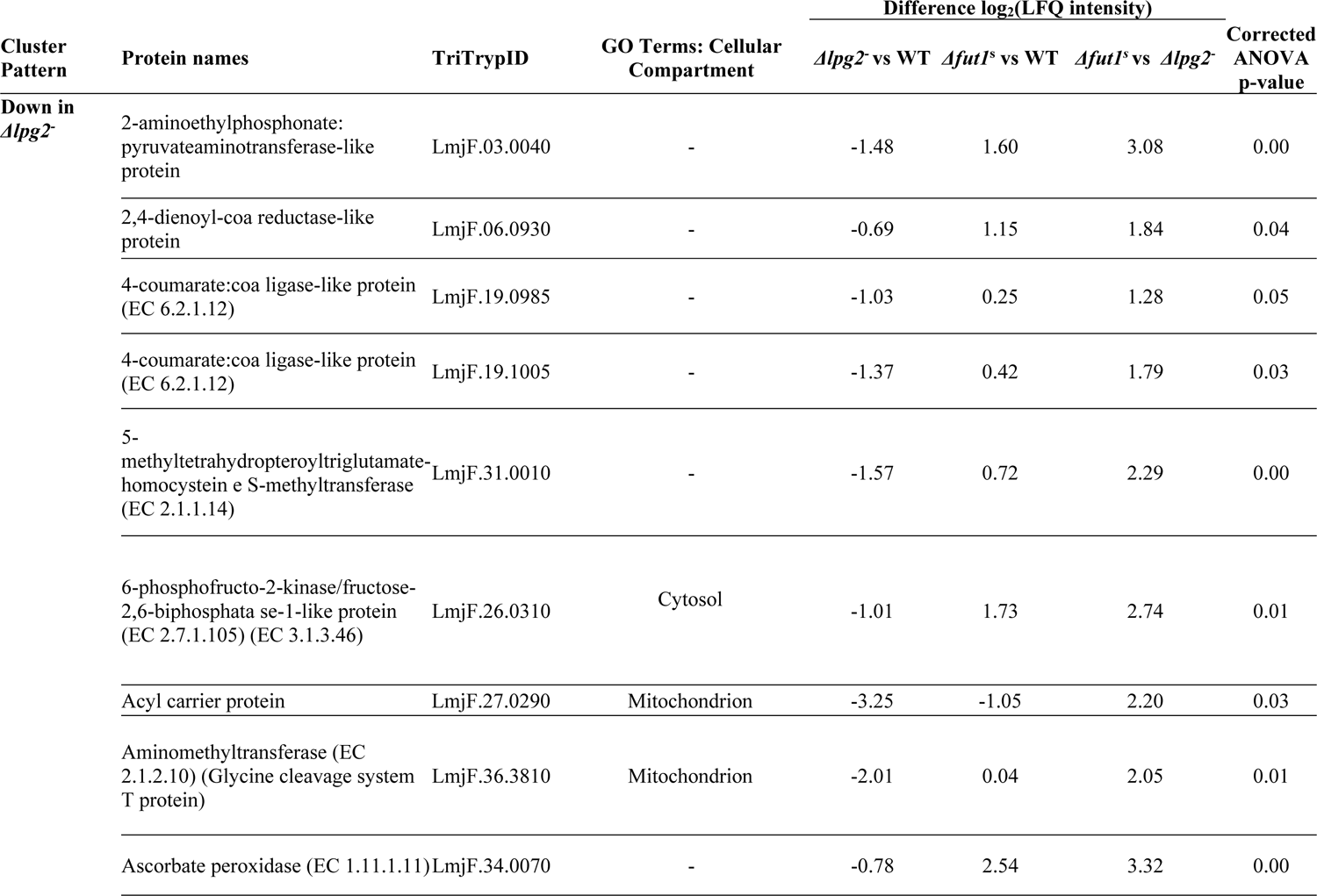
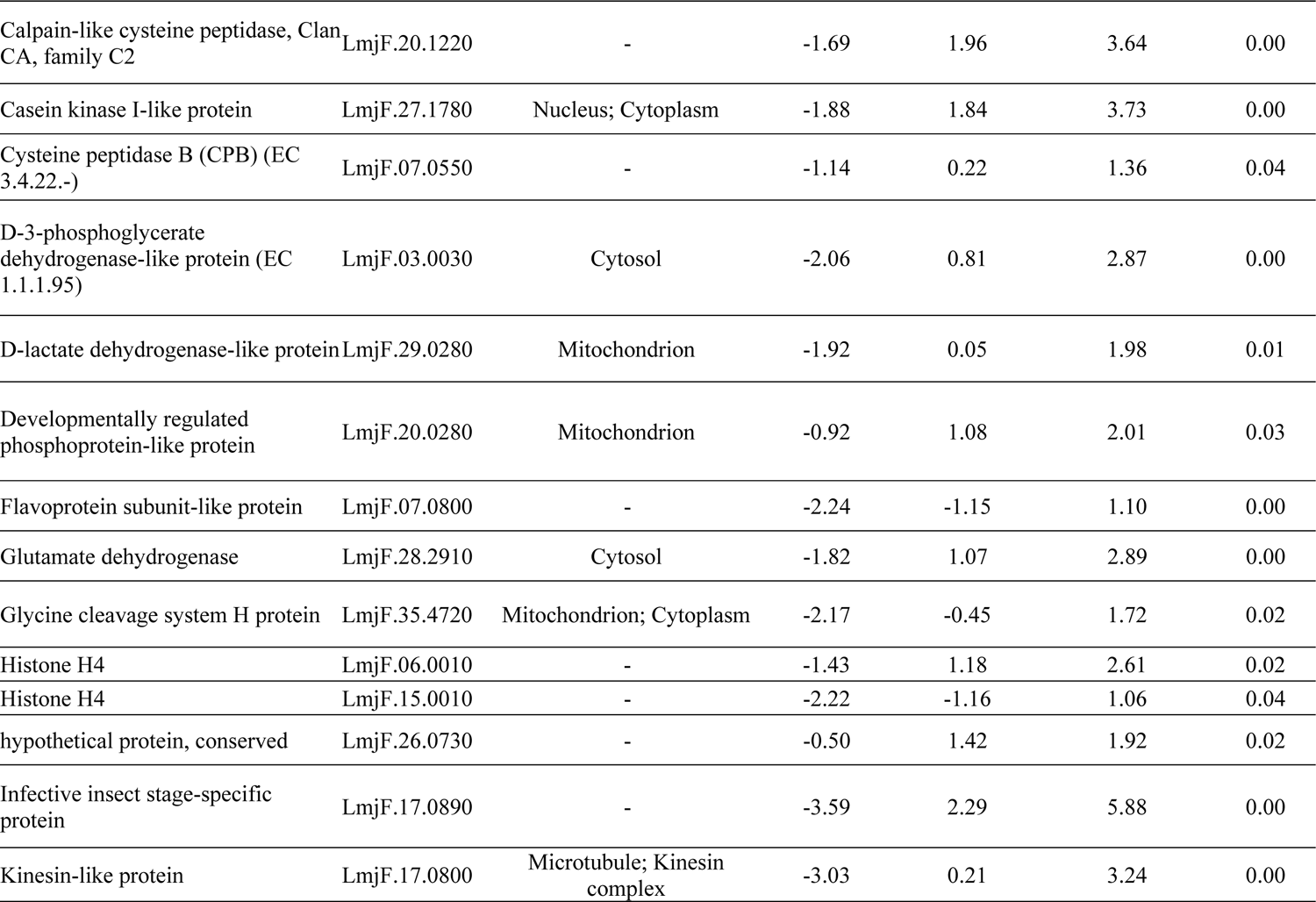
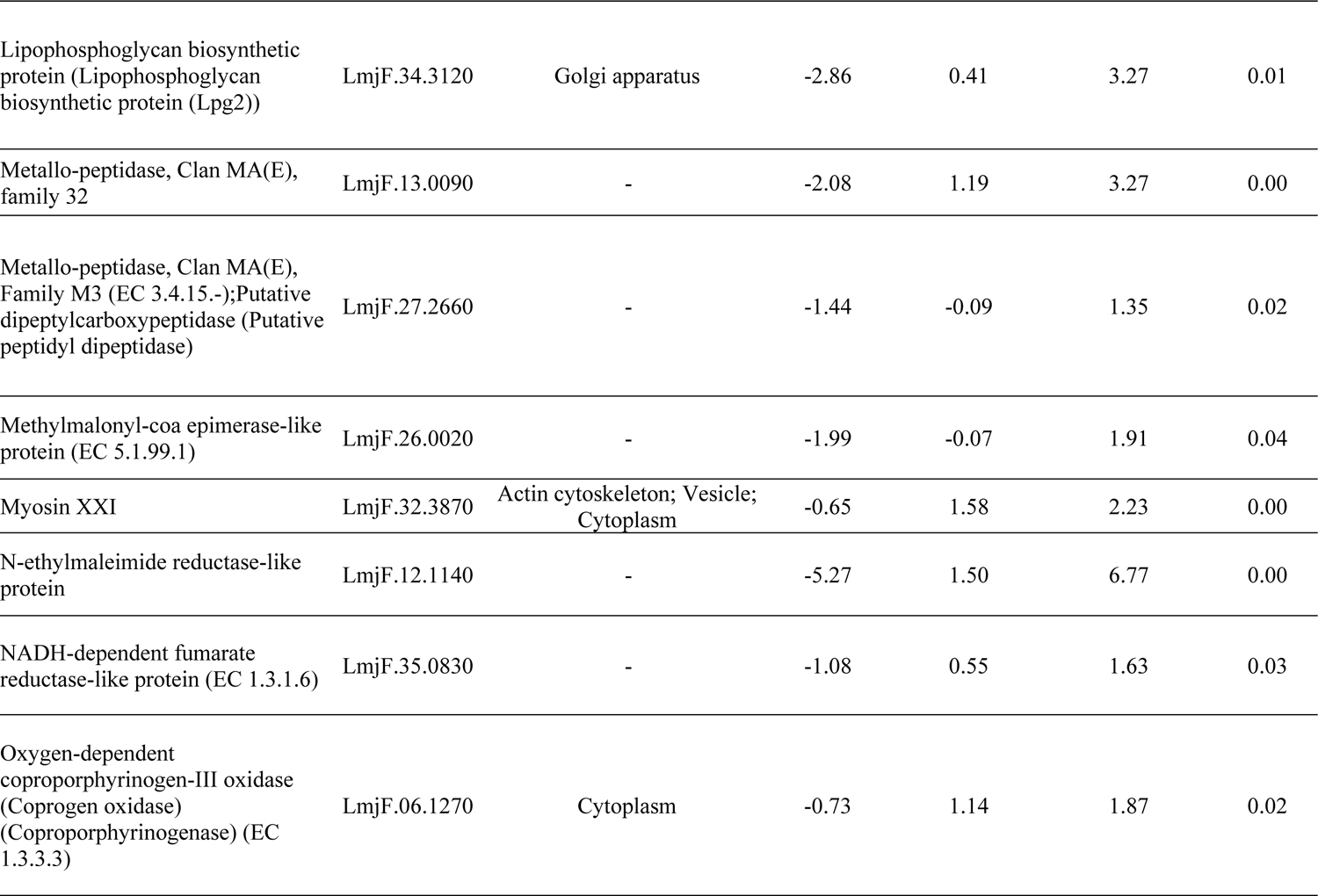
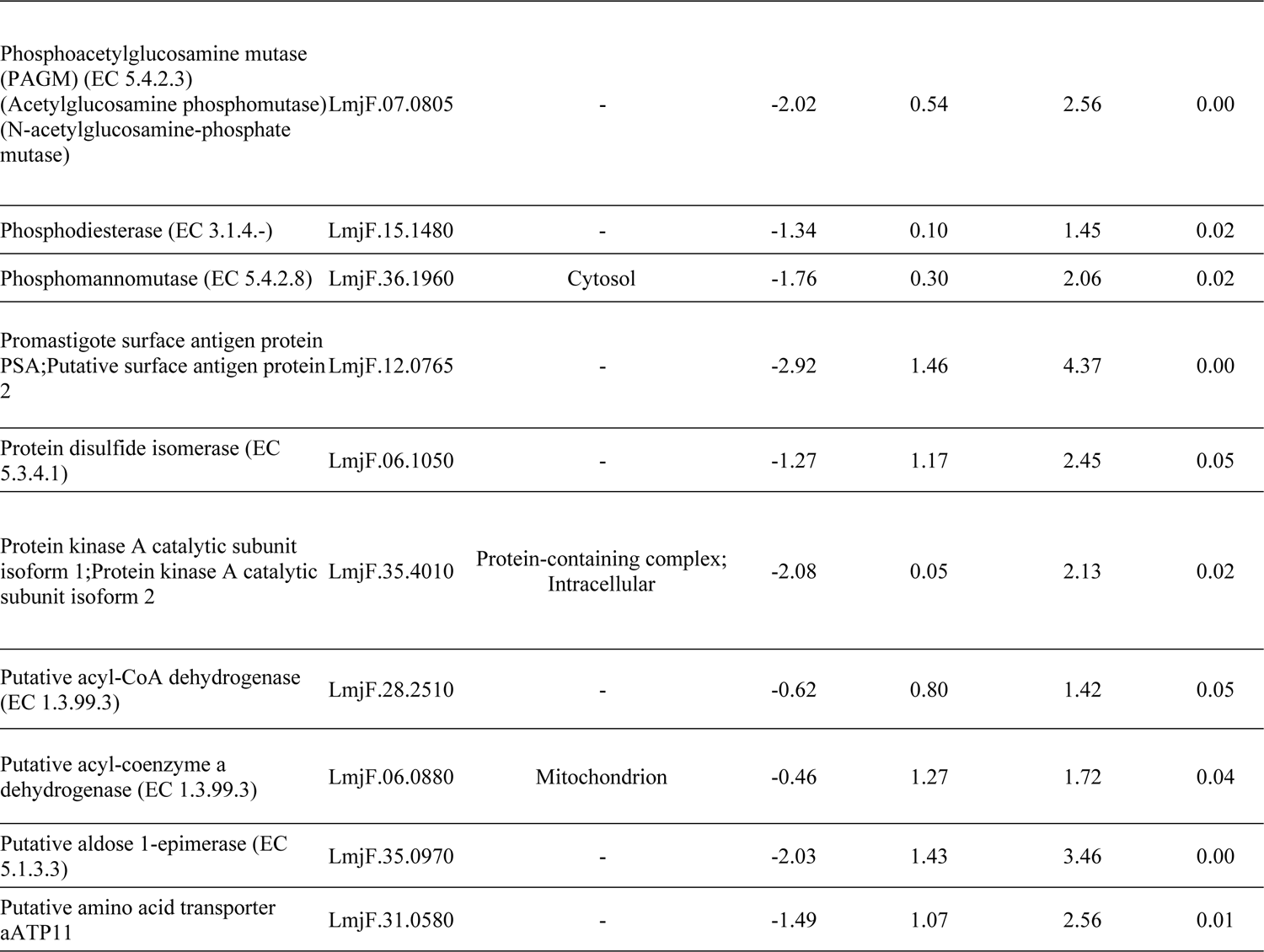
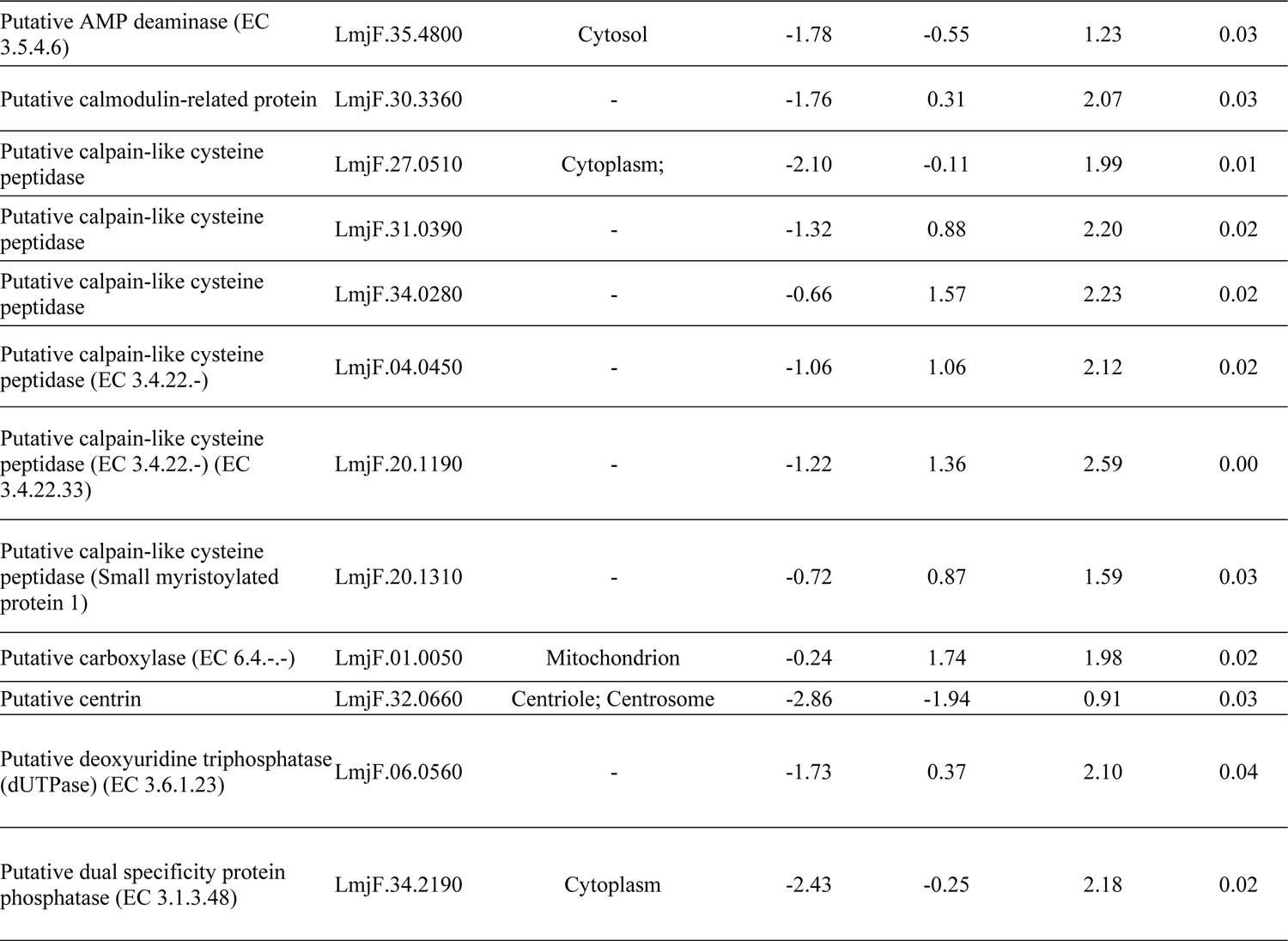
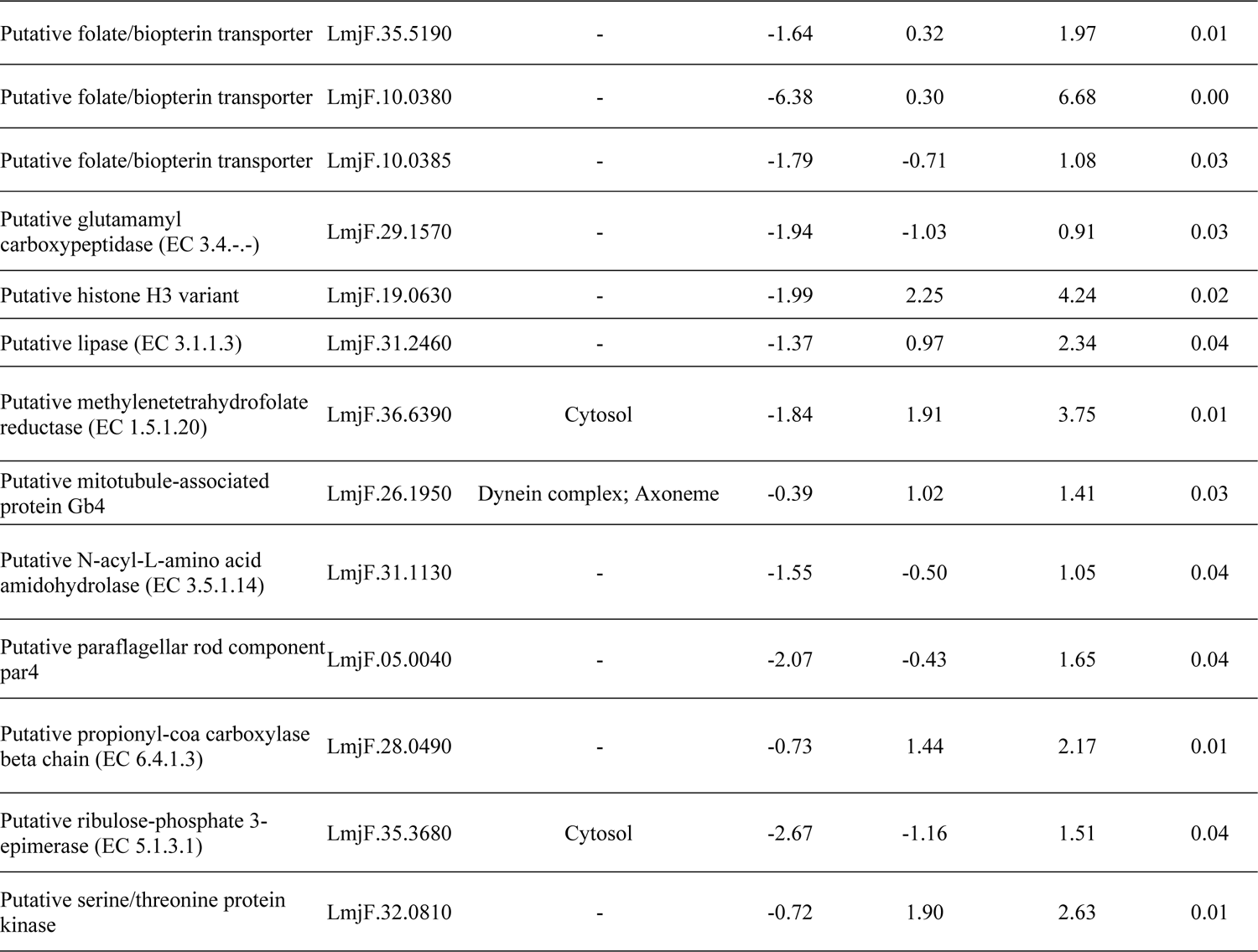
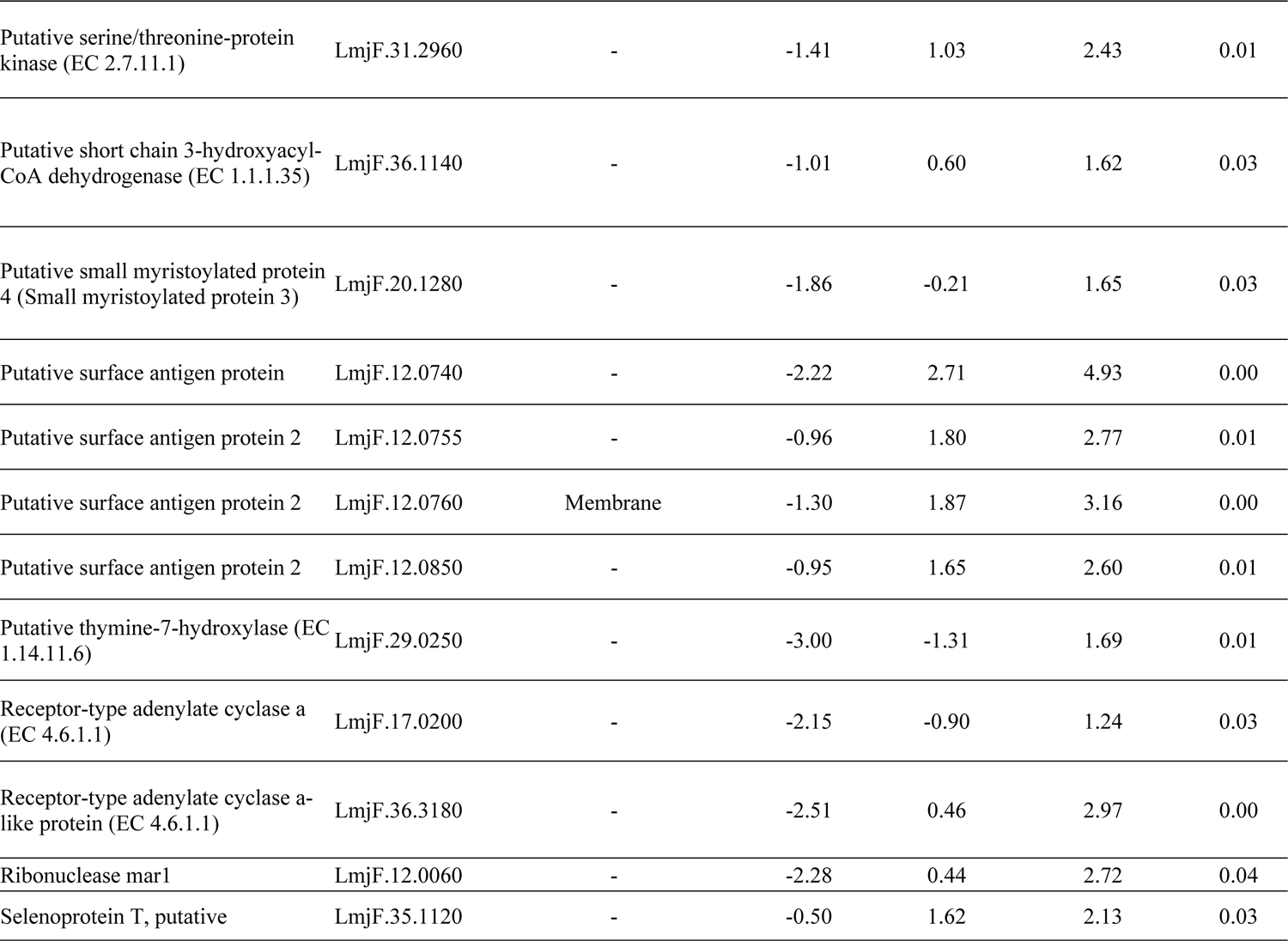
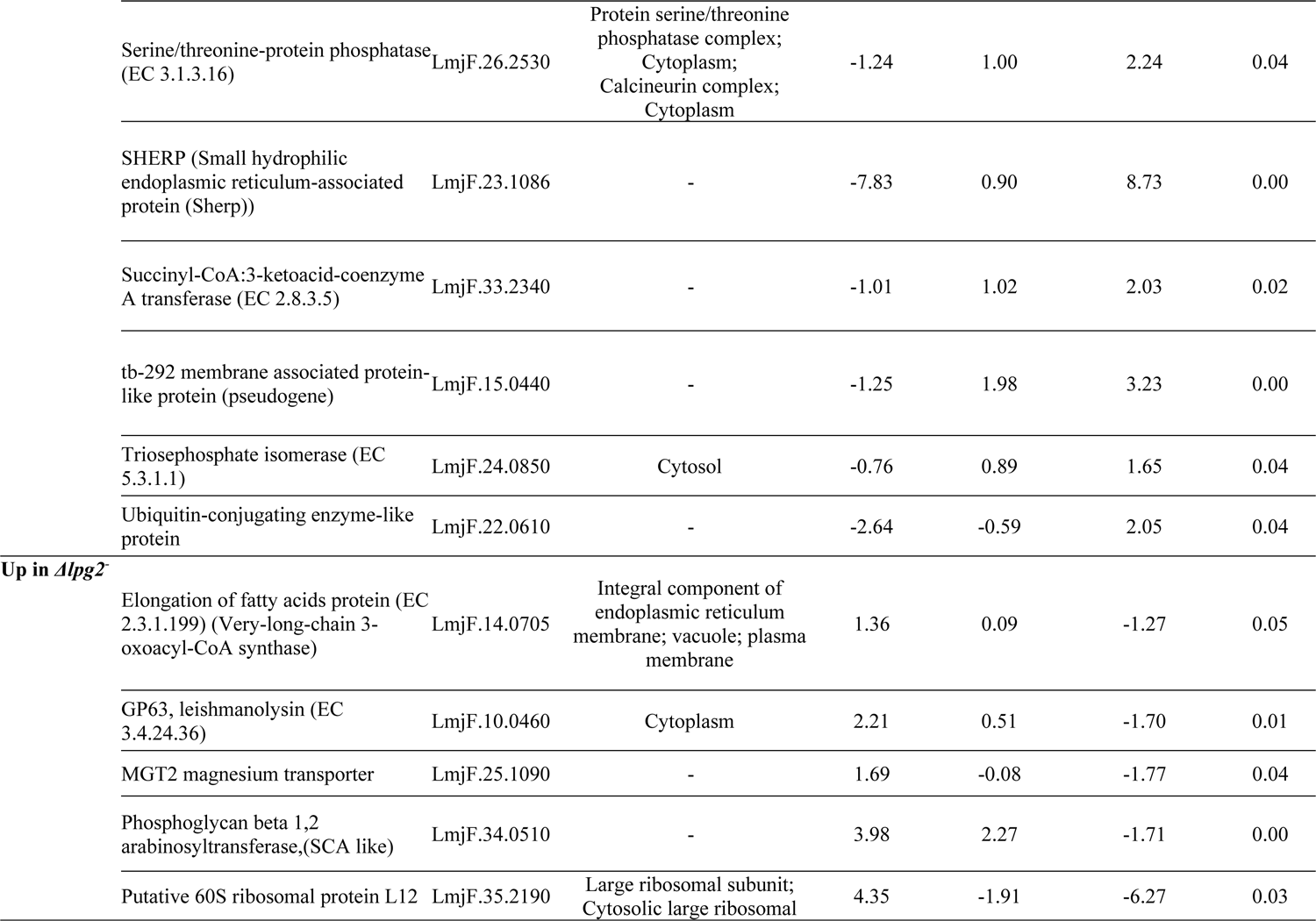
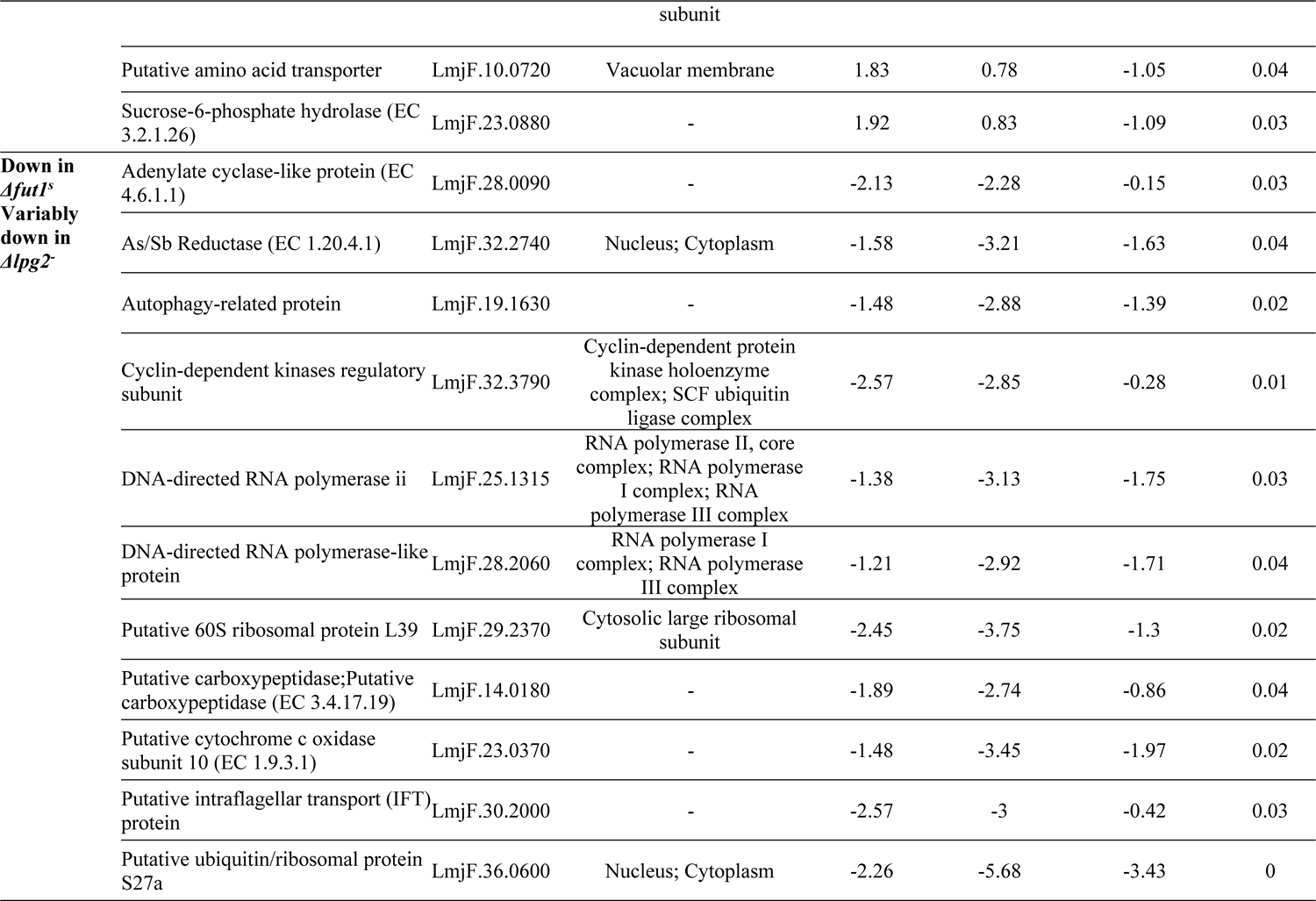
Proteins Significantly Affected in *Δlpg2^-^*. Proteins differing significantly in abundance between the *Δfut1**^s^**, Δlpg2^-^,* and WT parasite lines were evaluated by ANOVA; those proteins within clusters changing in *Δlpg2^-^*are shown (Fig. 4; Table S3). Overall, there were 174 proteins significantly decreased in *Δlpg2^-^*, of which 89 are of unknown function, while 13 increased in *Δlpg2^-^* of which 6 are of unknown function. A third group of 30 proteins, of which 19 are of unknown function, were significantly decreased in *Δfut1^s^* but varied among the biological replicates in *Δlpg2^-^* (Table S5).

